# Historic mine waste contains diverse microbial communities that reflect waste type and geochemistry

**DOI:** 10.1101/2025.02.25.640082

**Authors:** Mackenzie B. Best, Zohreh Kazemi Motlagh, Virginia T. McLemore, Daniel S. Jones

## Abstract

Waste rock and tailings left behind by historic mining operations can contain substantial critical mineral resources. However, over the decades and centuries since these deposits were emplaced, microbial communities developed that can catalyze rock weathering and elemental cycling, which could have impacted the economic resources but also might be harnessed for future biomining or other metal recovery efforts. Here we combined cell counting, rRNA gene and transcript sequencing, and whole rock geochemistry to compare the composition and abundance of microbial communities from five inactive mine sites in South-Central New Mexico that contain critical minerals. While acidic seeps and adits at the sites contained organisms commonly found in acid rock drainage and bioleaching operations, these organisms were only present at very low abundance in the waste rock and tailings, which were instead dominated by bacteria and archaea that mostly represent inorganic nitrogen- and organic carbon-oxidizing populations. Generally, rRNA transcript libraries contained many of the same organisms as rRNA gene libraries, indicating that most of these microbial populations are active. Differences among total and active communities correspond to waste rock geochemistry, including concentrations of sulfur, iron, and other variables such as copper, lead, and rare earth elements. Nevertheless, many of the rRNA gene and transcript sequences in these deposits were from groups without cultured representatives, and these unknown microorganisms are likely important for biogeochemical cycling over the long lifetime of these waste deposits. We also discuss recommendations for microbiological assessment of similar large historic mine waste deposits.

**IMPORTANCE:** New Mexico has a long history of mining, with hundreds of mining districts across the state, many of which contain inactive operations with historic tailings and waste rock. Because metallurgical processing was in its infancy when most of these mines were active, they contain substantial metal resources in tailings and waste rock that could be used to support domestic demand for critical minerals. We found that microbial communities associated with these deposits do not represent typical bioleaching communities, and instead are dominated by taxa not typically associated with mine wastes. However, the deposits do contain rare iron and sulfur-cycling taxa that could catalyze metal mobilization, as well as active populations of novel microorganisms that are likely important for biogeochemical cycling. These mine waste-associated microbial communities could represent important resources for bioremediation and other biotechnological applications to recover valuable elements from these and other historic mine waste deposits.

## INTRODUCTION

Many of the low-carbon technologies at the heart of energy-efficient technologies are more metal and mineral intensive than traditional fossil fuel-based infrastructure, resulting in a huge increase in global demand and price for these metals and mineral commodities (1). These non-fuel commodities have been termed ‘critical minerals’ by the U.S. Geological Survey (USGS) and Department of Energy (DOE) due to their importance in the development of new technologies and large-scale infrastructure projects (2–4). However, many of the high-grade shallow ore deposits we have historically relied on to supply these minerals are dwindling, with some element reserves projected to be completely exhausted within 50-100 years (5). The result is that this increase in demand must be met with deeper, lower grade deposits (6) and has prompted a search for non-traditional repositories of critical minerals such as recycling pre-existing metal resources (7, 8) and reevaluating mineral deposits and mine wastes (9–16). The reliance on imported critical minerals means the supply chain is highly susceptible to disruption due to import costs as well as geopolitical factors such as trade negotiations and agreements (17). Given this vulnerability, there is a push to locate and quantify critical minerals within the US at both active and inactive mine sites to better support domestic demand. New Mexico has a long history with mining and contains over 9000 inactive mines, many of which contain waste material in the form of bulk waste rock or tailings that were mined from precious- and base-metal deposits that have elevated concentrations of critical minerals (9, 18). These inactive mines vary widely in age and production history, and were processed exclusively for select base and precious metals at a time when metallurgical processing was in its infancy, meaning that they could contain substantial critical mineral resources within these wastes.

Microorganisms matter for each step in the mining lifecycle due to their capacity to enhance mineral weathering and mobilize and precipitate metals (19–23). When sulfide-rich minerals are exposed to the surface, microbially-catalyzed redox reactions can lead to the formation of acid rock drainage (ARD) or acid mine drainage (AMD) (24–35). Biological or abiotic strategies to prevent or limit sulfide mineral-oxidizing populations can help prevent AMD and secure mine wastes (36, 37), and acidophilic populations can be used to immobilize and even recover metals using active and passive remediation strategies such as low pH iron oxidation and anaerobic sulfate reduction (38–43).

Microorganisms can also be used to extract metals during active mining in ‘bioleaching’ or ‘biooxidation’ processes that are collectively known as “biomining” (44–50). The ability of microbes to oxidize metal sulfide and other minerals by producing oxidants, acids, and lixiviants, either during direct mineral contact or in a leaching solution (51–53) (54, 55), lowers the processing cost, which has made it easier for companies to process non-traditional metal resources such as refractory and low-grade ores (49, 52, 56, 57), electronic waste (58–60), and mine waste (61–63). Currently, approximately 15% of global copper, as well as smaller amounts of gold and other metals, is extracted using microorganisms (64, 65). Microbes have also been evaluated for in situ leaching concepts, in which the rock is first fractured and treated with a biologically-regenerated leach solution (e.g., (65–67)). Although mostly untested, mine wastes are potential targets for in situ bioleaching because it is often relatively permeable (68, 69) and contains leftover metals. Small-scale studies have shown that biomining process does work with tailings (61, 62).

Mine wastes can be defined as any product from a mine that is left over once mining has ceased. This can include bulk waste rock and tailings. Bulk waste rock ranges in size from pebble to boulder and includes materials that were above or within the ores (overburden and gangue, respectively) and was excavated and stored in waste piles on the surface. Bulk waste rock has not been processed for metal extraction, and may contain variable metal content (70–72). Tailings represent material that has been processed and metals extracted, leaving a fine-grained sand-sized residue. Although this material has been processed, many of these tailings still contain some metals (11–13, 73–77). Despite the importance of microbes for metal mobilization, relatively few studies have been carried out on these mine wastes to determine what microbial communities are present. From the limited number of studies addressing this knowledge gap, we know that these sources do contain robust communities of microorganisms, many of which can catalyze metal oxidation and reduction (23, 53, 73, 75, 78–87). These communities are highly variable and demonstrate a heterogeneity that is often overlooked when remediation strategies and modeling metal removal and cycling dynamics (23, 74), as these communities vary not just with metal availability, but with depth (83, 86, 88, 89). However, many of these studies have relied on traditional culture-based microbiological techniques, introducing bias by selecting for organisms that are easier to grow in lab settings and do not provide an accurate representation for environmental communities (73, 78, 81, 85). Understanding microbial community structure and variability across these types of mine waste is crucial to understanding metal cycling dynamics for these mining-impacted environments and may play a role in assessing the bioleaching potential of these low-grade non-traditional metal resources.

In order to evaluate the microbiological communities associated with historic mine wastes, we sampled waste rock, tailings, and sediment from five inactive mines in south-central New Mexico: Copper Flat, Center, Carlisle, Alhambra, and Black Hawk. This work was paired with geochemical studies assessing critical minerals in these wastes (90, 91). All five sites experienced production at some point in the past, and all contain critical minerals in their waste (90, 91). The goals of this study were to (i) evaluate the drivers behind the composition and structure of microbial communities in different waste types; (ii) evaluate methods for culture-independent analysis of historic mine wastes, including approaches for obtaining representative microbiological samples of large waste piles; and (iii) explore microbial resources for future in situ leaching opportunities and potential bioremediation strategies for these and other historical mining areas.

### Geologic context and mine waste characteristics

Samples for this study come from five inactive mines in three different mining districts: the Carlisle and Center mines in the Steeple Rock district, the Copper Flat mine in the Hillsboro district, and the Black Hawk and Alhambra mines in the Black Hawk district (Fig. 1).

**FIG 1.**
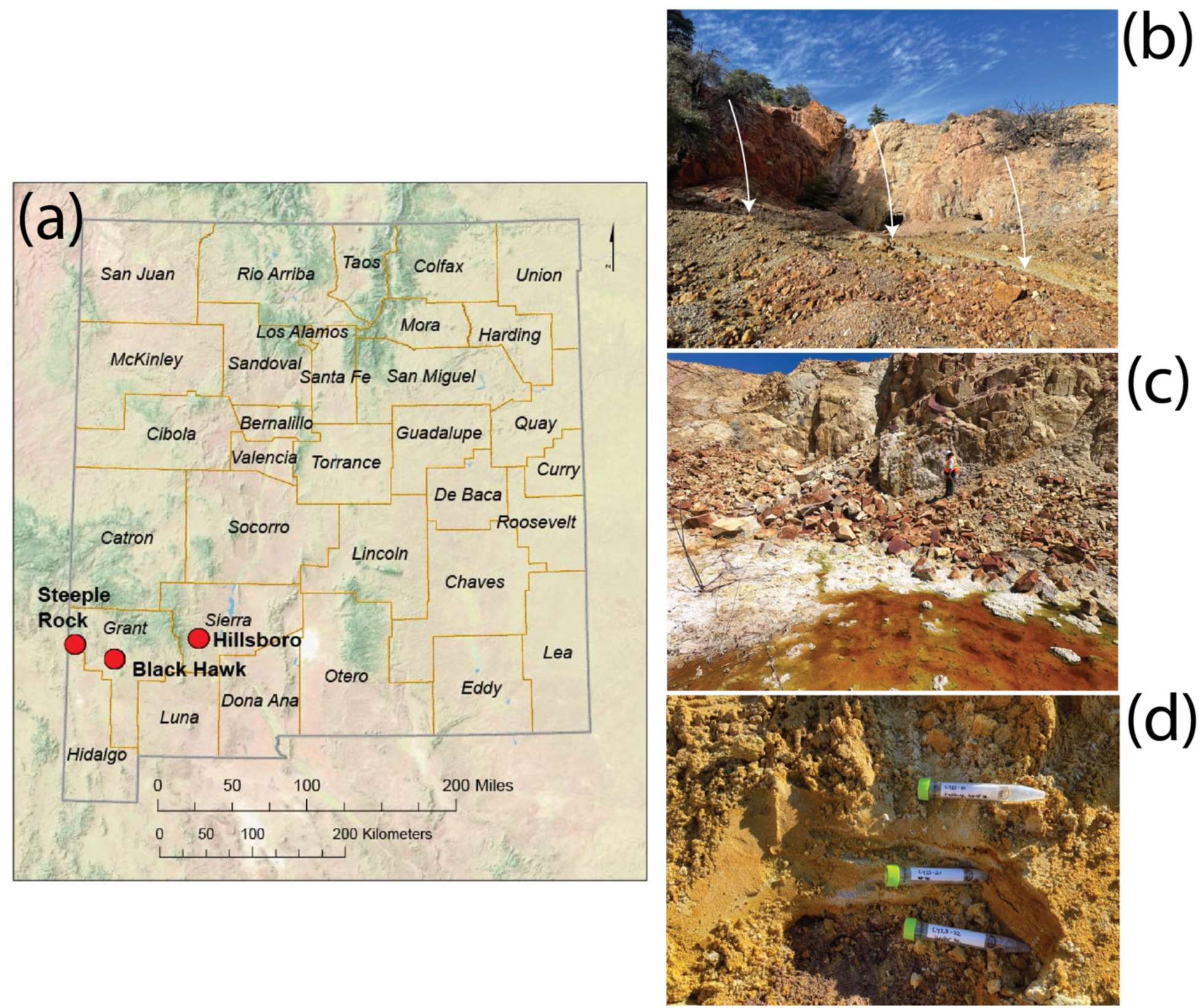
(a) Map of New Mexico showing the three districts sampled. The Hillsboro district contains the Copper Flat mine (CF), the Steeple Rock district contains the Center (CN) and Carlisle mines (CY), and the Black Hawk District contains the Black Hawk (BH) and Alhambra mines (AH). Representative photos show (b) bulk waste rock (white arrows) at CY, (c) a seasonal acidic seep at CF, and (d) tailings at CY.

The deposits at the Center and Carlisle mines are characterized by a low-sulfidation volcanic epithermal system with Au-Ag veins (92, 93). Exploration began in 1860, and production was first reported in 1880 and continued sporadically until 1994. An estimated $10 million worth of metals was produced including gold (Au), silver (Ag), copper (Cu), lead (Pb), and zinc (Zn) in its 100 years of activity (92). The workings at the Center mine includes a central flooded adit, and a mix of low-grade ore material and bulk waste rock (gangue and low-grade mineralization) that was piled up, flattened, and covered with 15 cm of soil cover. Recent work by (90) has revealed the presence of critical minerals in the mine waste at these sites, including arsenic (As), bismuth (Bi), tellurium (Te), fluorite (F), and Zn, all of which are associated with these types of mineral deposits.

The Copper Flat mine in the Hillsboro District was developed around the mineralized Copper Flat quartz-monzonite porphyry (CFQM) deposit that was first identified in 1975 (94, 95). Despite being in production for only 4 months in 1982, the Copper Flat mine produced 1.2 million short tons of ore containing Cu, Au, and Ag (96). This site features an open pit lake, a seasonal acidic seep, and extensive bulk waste rock piles and tailings deposits that contain substantial critical minerals endowments of Bi, cobalt (Co), gallium (Ga), and Te (90).

The Black Hawk mine is an arsenide 5-element vein mineral system. This deposit type is associated with critical minerals such as Co, nickel (Ni), As, Bi, tungsten (W), Te, and Zn. The site experienced sporadic production from 1881-1960, producing Cu, Au, and Pb (97). The nearby Alhambra mine also experienced production starting in 1881 (98). It is estimated that the district produced more than $1,000,000 silver from 1883 to 1893 (98). Recent geochemical work by (91) has revealed the presence of critical minerals in the mine wastes at both sites including cadmium (Cd), Zn, and Te. The mine waste at both inactive mines is in the form of bulk waste rock piles mixed with ore material.

## RESULTS

### Field sampling and observations

We collected samples from all five inactive mine sites: the Carlisle and Center mines in the Steeple Rock District (CY and CN, respectively), the Copper Flat mine in the Hillsboro district (CF), and the Black Hawk and Alhambra mines in the Black Hawk District (BH and AH, respectively) (Fig. 1). The waste types sampled at each location are summarized in Table 1, with additional details in Table S1.

**TABLE 1.** Waste rock types sampled at inactive mine sites.

| Mine | Bulk waste rock | Acidic seep | Flooded adit | Tailings |
| --- | --- | --- | --- | --- |
| Copper Flat (CF) | Y | Y | - | Y |
| Center (CN) | Y | - | Y | - |
| Carlisle (CY) | Y | - | Y | Y |
| Black Hawk (BH) | Y | - | - | - |
| Alhambra (AH) | Y | - | - | - |

“-” indicates this sample type was not present (and therefore not collected)

Bulk waste rock was collected from all five sites. In order to take a representative sample of a large waste rock pile, a grid of approximately equal-sized squares was superimposed onto the waste rock pile and the center of each grid square demarcated by flags. Individual samples are collected at each flag (following protocols established by the USGS, outlined in (9). The overburden was removed, and the sample was sieved through a 4mm sieve for subsequent geochemical analyses. To get a representative composite sample for the entirety of the waste rock pile, a small but equal volume of material from each individual sample hole was sieved through a 4mm sieve and combined into one composite sample. For microbiological sampling, six individual samples were chosen, and samples collected from each individual hole. Similarly, a composite sample was collected by combining small amounts of material from each of the selected individual geochemistry holes. Geochemical data from these sites were reported in (91) and (90).

Tailings were sampled at CF and CY. The tailings at CF are horizontally and vertically extensive, and so we employed the same sampling strategy as with the bulk waste rock piles: six individual samples and a composite sample were collected. In addition, tailings were collected above, at, and below a visible oxidation front from one deep sampling pit (Fig. S1). The distance from the top of the pile to the oxidation front was 30 cm, and the distance from the oxidation front to the bottom of the hole where the third sample was collected was 70 cm. The tailings at CY are much less extensive both horizontally and vertically, so only three samples were collected: above, at, and below a visible oxidation interface like at CF (Fig. 1a).

Sediment and biofilms were sampled from flooded adits at CN and CY. At the time of sampling, the pH of the CN adit was pH 6.5, while the two flooded adits at CY were pH 3.1 and 3.38 (Table S1). Sediments were also sampled from the seasonal acidic seep at CF where the pH measured 3.55 in 2022 and between 1.54 and 1.95 in 2020.

### 16S rRNA gene libraries

We prepared 93 successful rRNA gene libraries from 64 of 73 total samples attempted. Libraries ranged from 10,094 to 104,001 reads per sample with an average library size of 43,269 reads (standard deviation 24,048). In total, 15,648 operating taxonomic units (OTUs) were identified at 97% similarity.

The three types of waste were associated with different microbial communities. The acidic seep samples from CF and the flooded adits of CN and CY were dominated by genera commonly found in ARD such as *Acidiphilium, Leptospirillum, Acidibacter, Metallibacterium,* and *Ferrithrix* (Fig. S2). In contrast, the waste rock samples from all sites show much more diverse communities compared to the seeps and tailings (Fig. 2), and include abundant bacteria in the genera *Bradyrhizobium* and *Solirubrobacter*, along with abundant archaea such as *Candidatus Nitrocosmicus* and *Nitrososphaeraceae* spp. (Fig. S3). The tailings also contain some of these same genera, as well as *Ramlibacter*, *Paucibacter*, *Enhydrobacter*, *Kocuria*, unclassified *Ktedonobacteraceae*, and *Spingomonas*. Some of the genera associated with iron and sulfur oxidation that occur in the seep and adit sediments are present at low levels in other waste types, such as *Acidiphilium* (Fig. 3), which also contain some other rare taxa associated with metal and sulfur oxidation or reduction such as *Sulfuriferula*, *Acidibacter,* and *Sulfurifustis* (Fig. 3). Overall, the microbial communities from both waste rock and tailings are dominated at all sites by uncultured and sometimes unclassified microorganisms (Fig. 3, S4).

**FIG 2.**
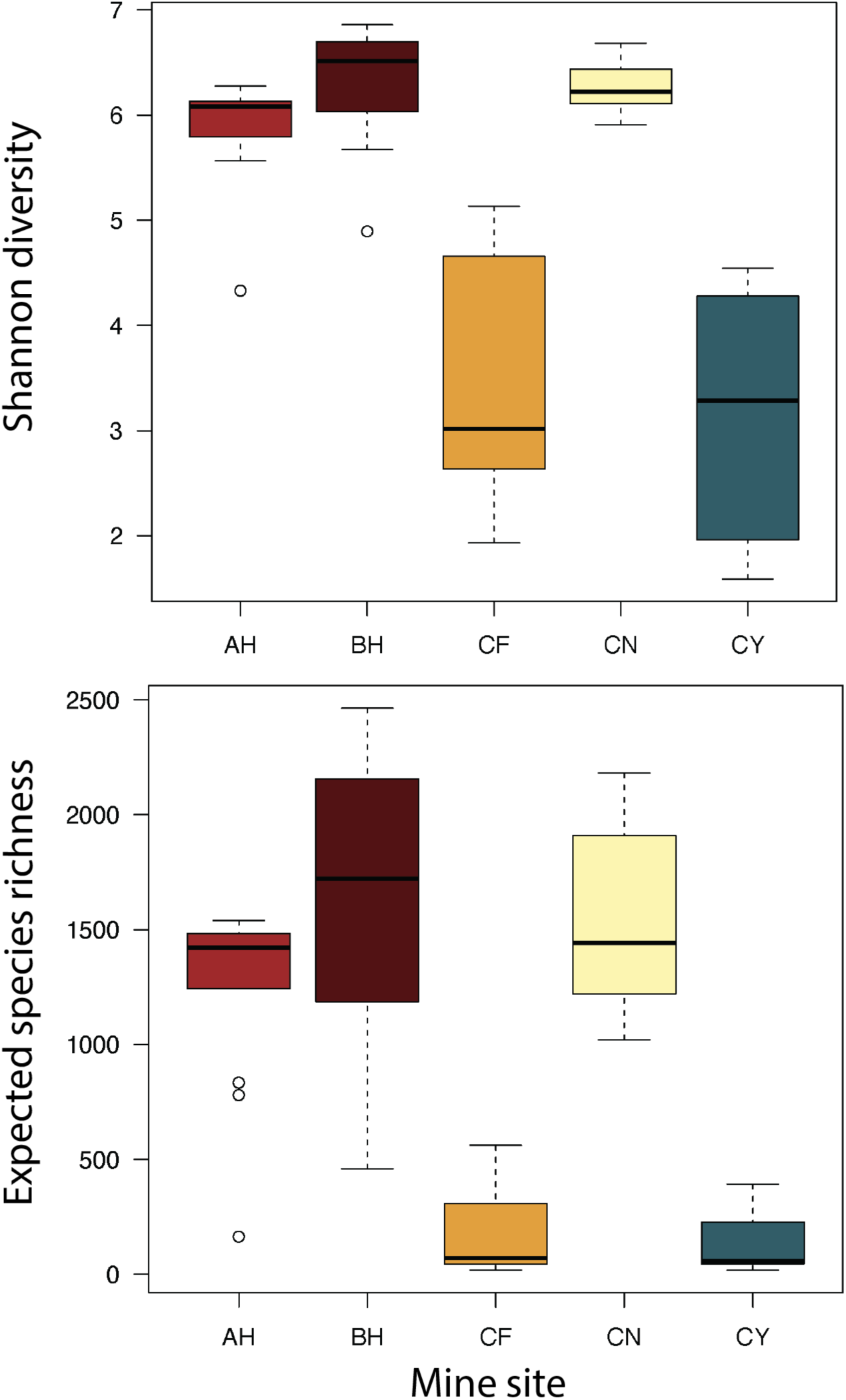
Shannon diversity (top) and expected species richness (bottom) boxplots of bulk waste rock and tailings samples from the Alhambra mine (AH), Black Hawk mine (BH), Copper Flat mine (CF), Center mine (CN), and Carlisle mine (CY). Sites CF and CY did not have a soil cover.

**FIG 3.**
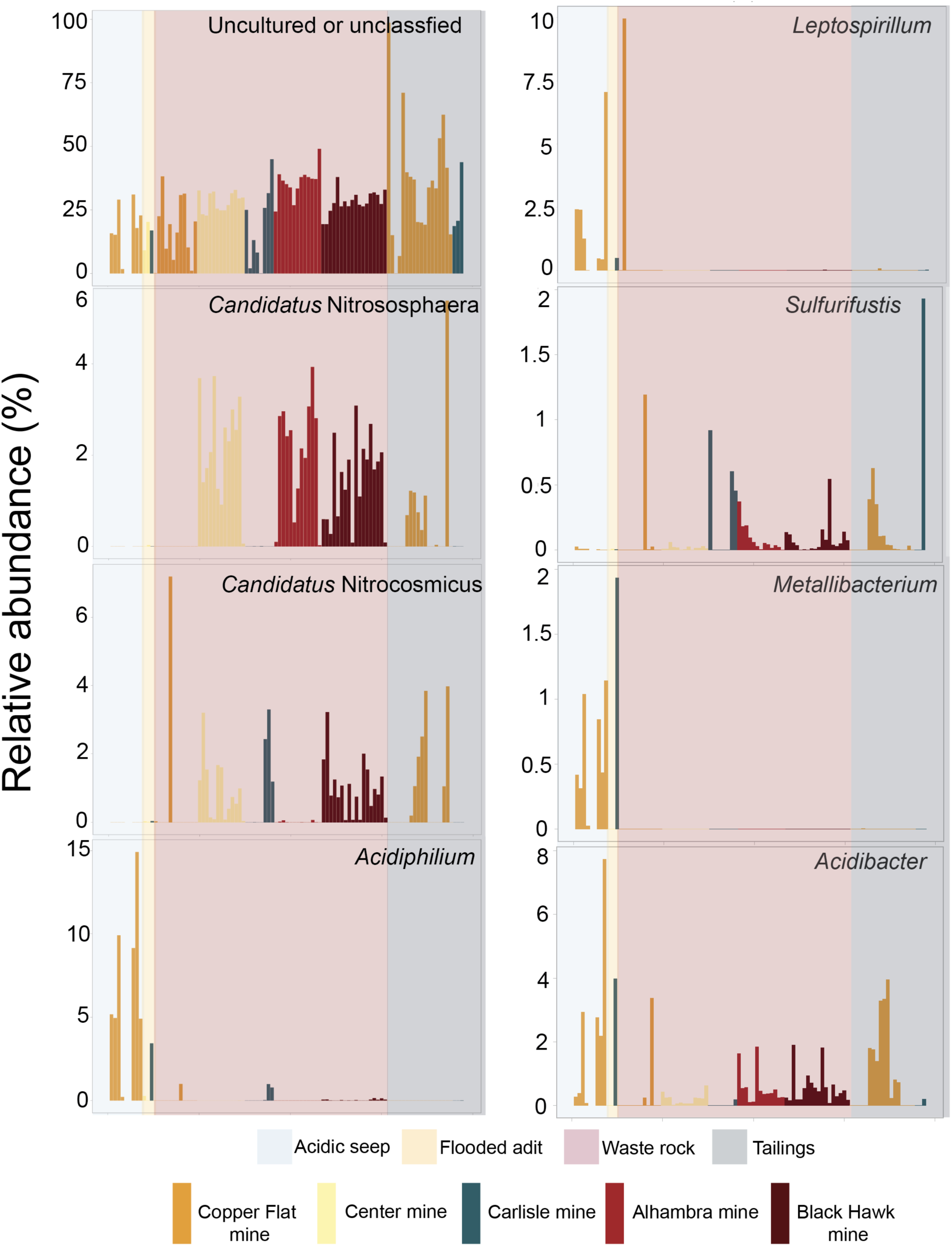
Bar chart showing the relative abundance of specific taxa in waste samples by sample type and location. ‘Unclassified’ or ‘uncultured’ refers to OTUs that were either unclassified at the genus level or higher, or from groups with no known cultured representative.

Non-metric multidimensional scaling (NMDS) ordination shows that rRNA gene libraries from sites AH, BH, and CN are similar to each other and cluster tightly at one extreme of the ordination, while libraries from sites CF and CY were much more heterogeneous (Fig. 4). Sites AH, BH, and CN also had the highest diversity (Fig. 2), perhaps because these are the three sites that had a soil cover. ANalysis Of SIMilarities (ANOSIM) analysis showed that libraries from different sites were statistically significantly different (R = 0.28, p < 0.001), as were libraries from bulk waste rock versus tailings tailings, for all sites (R = 0.28, p < 0.001) as well as for only libraries from sites CY and CF for which both tailings and waste rock were sampled (R = 0.40, p < 0.001). These differences were also statistically significant by PERMANOVA (by site, PERMANOVA F = 6.8, p < 0.001; waste rock versus tailings for all sites, PERMANOVA F = 6.0, p < 0.001; waste rock versus tailings for site CY and CF, PERMANOVA F = 3.6, p < 0.001), and all pairwise comparisons of libraries by site are also all statistically significant (PERMANOVA F = 2.3 - 8.7, p < 0.002).

**FIG 4.**
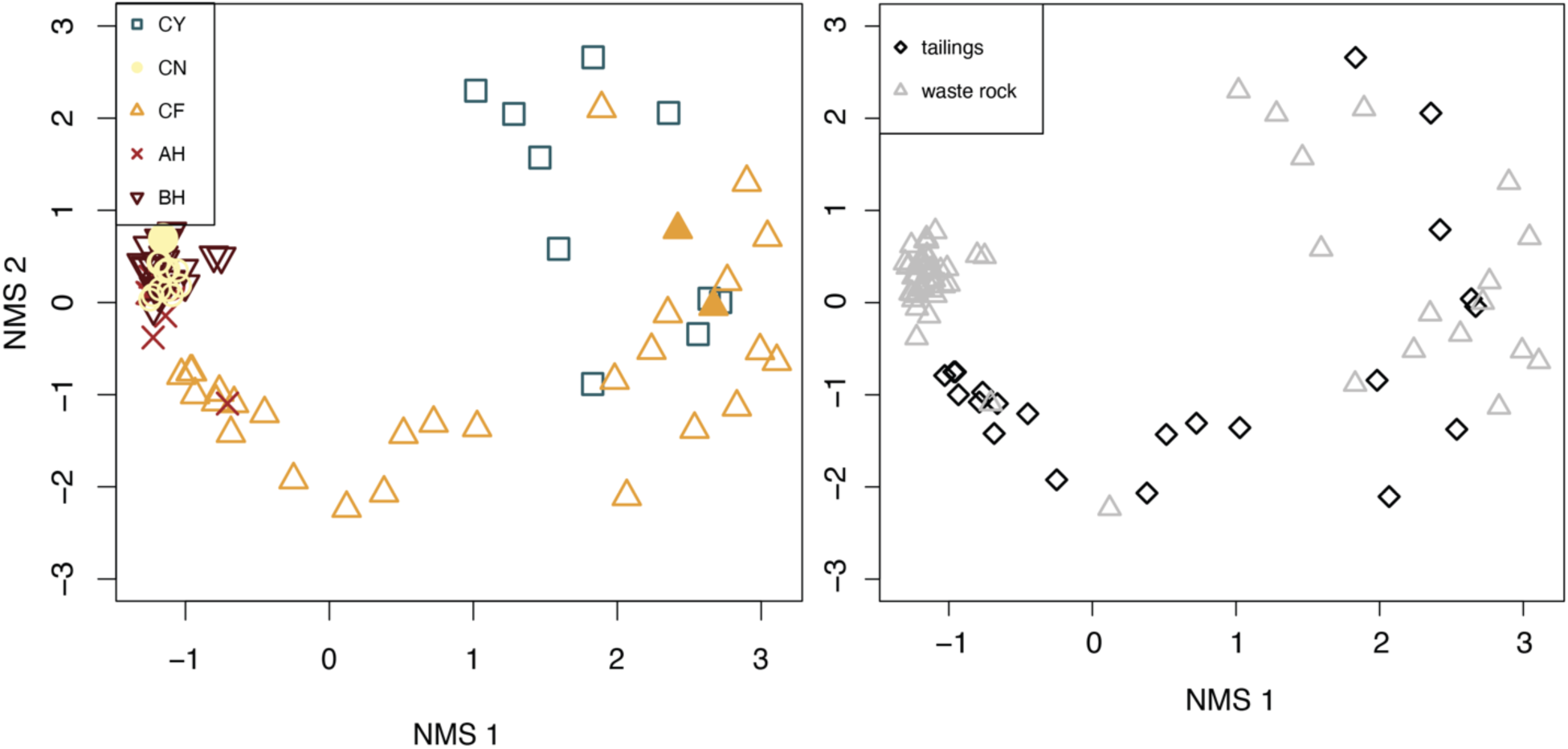
A non-metric multidimensional scaling (NMDS) ordination of samples collected from waste rock and tailings all five inactive mine sites (seep and adit sediments are not included). In the left panel, samples are coded by sample location. Unfilled shapes are rRNA gene libraries, and filled shapes are rRNA transcript libraries from composite samples. In the right panel, samples are coded by sample type (either waste rock or tailings).

In order to evaluate potential geochemical drivers of microbial community composition, we used an environmental overlay to evaluate the relationships among geochemical variables and rRNA gene and transcript libraries in NMDS ordinations (Fig. 5). When libraries from all sites are included, %S, %C, SiO_2_, and copper content are statistically significant and oriented with the first ordination axis. This aligns with the separation between libraries from CN, AH, and BH, which cluster tightly at the extreme of the first ordination axis, and libraries CY and CF, which are more variable. When only libraries from only sites AH, BH, and CN were ordinated, libraries cluster by site, with %S, As, Pb, and U oriented with the first ordination axis, and Cu, Co, Ag, and TREE (total rare earth element) concentration oriented positively with the first ordination axis and negatively with the second. When sites were compared based on geochemical variables alone, sites AH and BH overlap, and separate from CF and CY along the first ordination axis, indicating distinct geochemical characteristics that could be driving differences in microbial communities among the sites (Fig. S5). In order to further explore the relationship between specific taxa and geochemical variables, we correlated the most abundant OTUs with individual geochemical variables in order to determine if specific populations might be responding to specific geochemical drivers. Select correlations are shown in Fig. S6.

**FIG 5.**
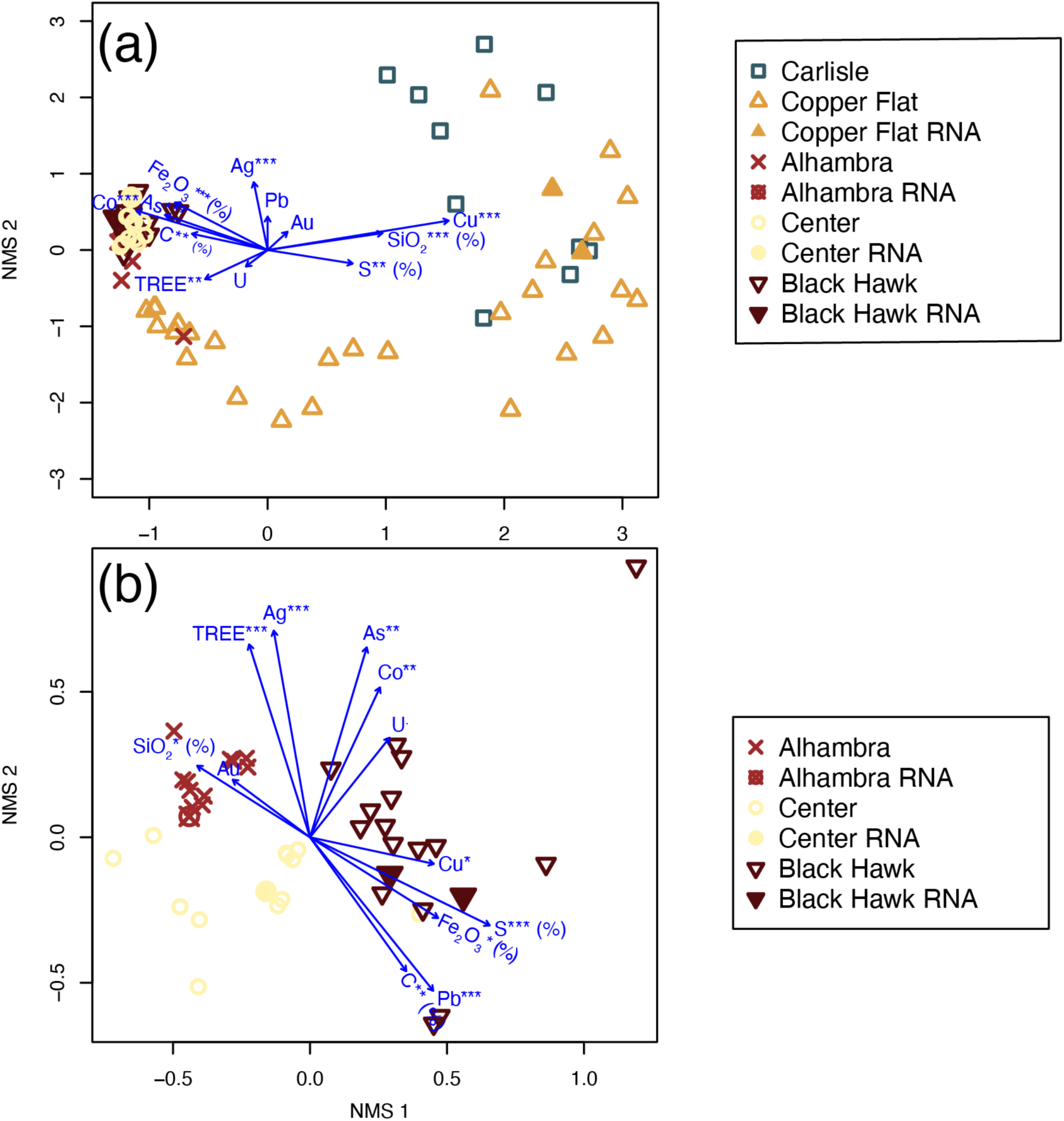
Non-metric multidimensional scaling (NMDS) ordinations of samples from (a) the Alhambra, Black Hawk, Copper Flat, Carlisle, and Center mines (AH, BH, CF, CY, and CN, respectively) and (b) only AH, BH, and CN, for which bulk waste rock was topped with a soil cover. Vectors of fitted geochemical variables indicate by asterisks are statistically significant (*** = p <0.001, ** = p <0.01, and * = p <0.05). RNA transcript samples are represented by filled shapes, and DNA samples are represented by open shapes.

### Evaluation of sampling methods for historic waste rock and tailings piles

To attempt to get a representative sample of large waste rock and tailings piles, six samples were taken from separate random locations around the perimeter and surface of the pile. A small volume of material from each of these individual samples were combined into a composite sample and homogenized, and libraries prepared from both the individual and composite samples. Non-metric multidimensional scaling (NMDS) ordination shows that the composite samples plot with the individual samples from the same waste pile for all five sites (Fig. 6). In addition, because waste rock is heterogeneous, we extracted each samples twice and, in some cases, sequenced individual sample DNA extracts along with the combined DNA extracts. Overall, 16S rRNA gene libraries from the individual extracts were similar to those from the combined extract (Fig. S7). We also found that the Qiagen PowerSoil Pro kit resulted in higher DNA yields than other kits tested, so all DNA library analyses reported in this study were extracted using this kit using a bead-beating protocol described in (99).

**FIG 6.**
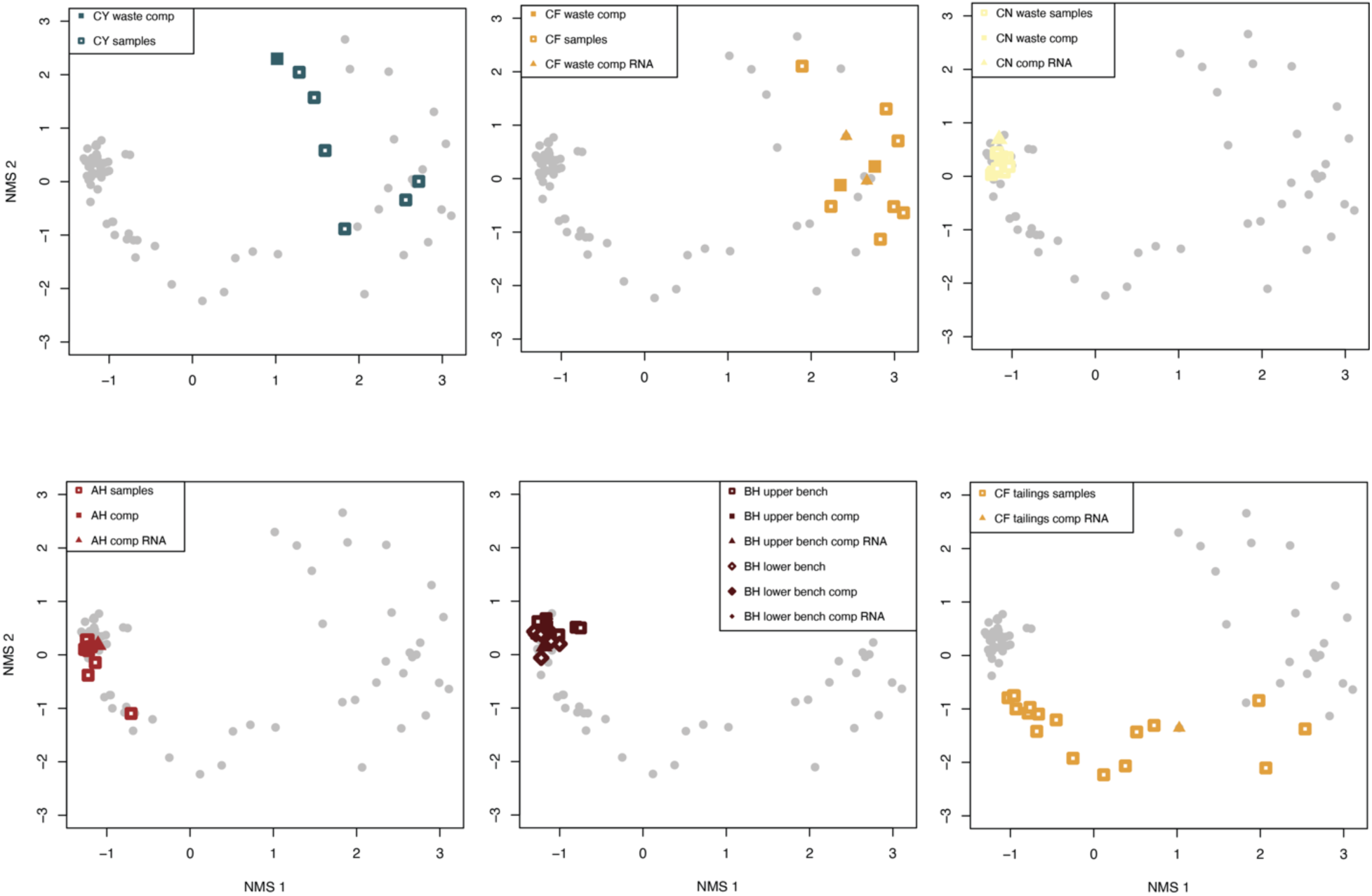
Non-metric multidimensional scaling (NMDS) ordinations of samples from each mine site comparing composite samples (filled symbols) and individual samples (open symbols) from the same locations. Squares indicate rRNA gene libraries, and triangles indicate rRNA transcript libraries. The bottom right panel is tailings from site CF, and the others are all waste rock. In the bottom middle panel, the upper and lower bench at site BH refers to samples collected from two distinct benches from previous mining activity. The upper bench refers to the bench that is higher in elevation (by 5m) that is the ‘top’ of the waste rock piles. In the legends, “comp” = composite sample.

### 16S rRNA transcript libraries

In order to assess active microbial populations in the mine waste, rRNA transcript libraries were prepared from composite samples of all waste rock piles and the CF tailings. rRNA transcript libraries from all except for one sample (CY23-13) were successful, and successful libraries contained between 34,975 and 57,123 with an average library size of 38,759 reads (standard deviation 8,140).

NMDS ordinations show that 16S rRNA transcript libraries from the composite waste rock, as well as the tailings composite from CF, are overall similar to 16S rRNA gene libraries from those locations (Fig. 4, 7, 8); NMDS analyses of rRNA gene libraries alone have similar structure to those with rRNA transcripts (Fig. S7). Of the seven composite samples with successful 16S rRNA transcript libraries, six had successful 16S rRNA gene libraries and could be compared directly. For the other composite tailings sample, rRNA transcript libraries were compared to individual samples from the same tailings pile. These comparisons show that, overall, most taxa in the rRNA gene libraries are present in the rRNA transcript libraries but there are individual OTUs and phyla that be either over-or under-represented (Fig. 7, 8). In general, archaea such as *Candidatus* Nitrososphaera and *Candidatus* Nitrocosmicus are represented at much higher abundances in the rRNA gene libraries compared to the rRNA transcript libraries. Other genera, such as *Bradyrhizobium* and *Streptomyces*, appear in higher abundance in the rRNA transcript libraries compared to the rRNA gene libraries. The majority of organisms appear similarly abundant in rRNA transcripts as in the gene libraries, such as *Solirubrobacter* spp., *Sulfurifustis* spp., *Nitrospira* spp., and *Kocuria* spp.

**FIG 7.**
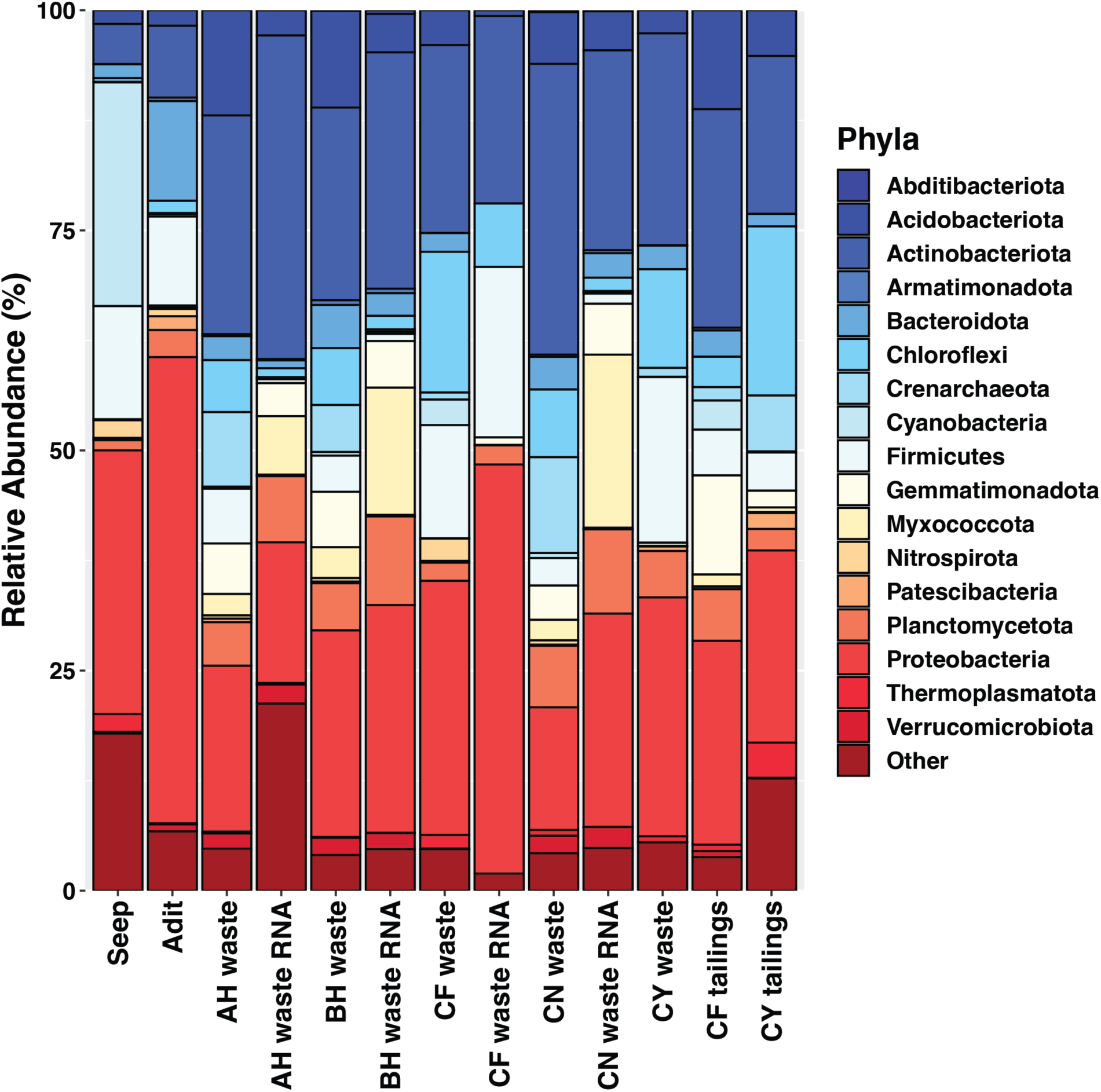
Phyla-level community composition of samples by sample type and rRNA gene versus rRNA transcript libraries from composite samples.

**FIG 8.**
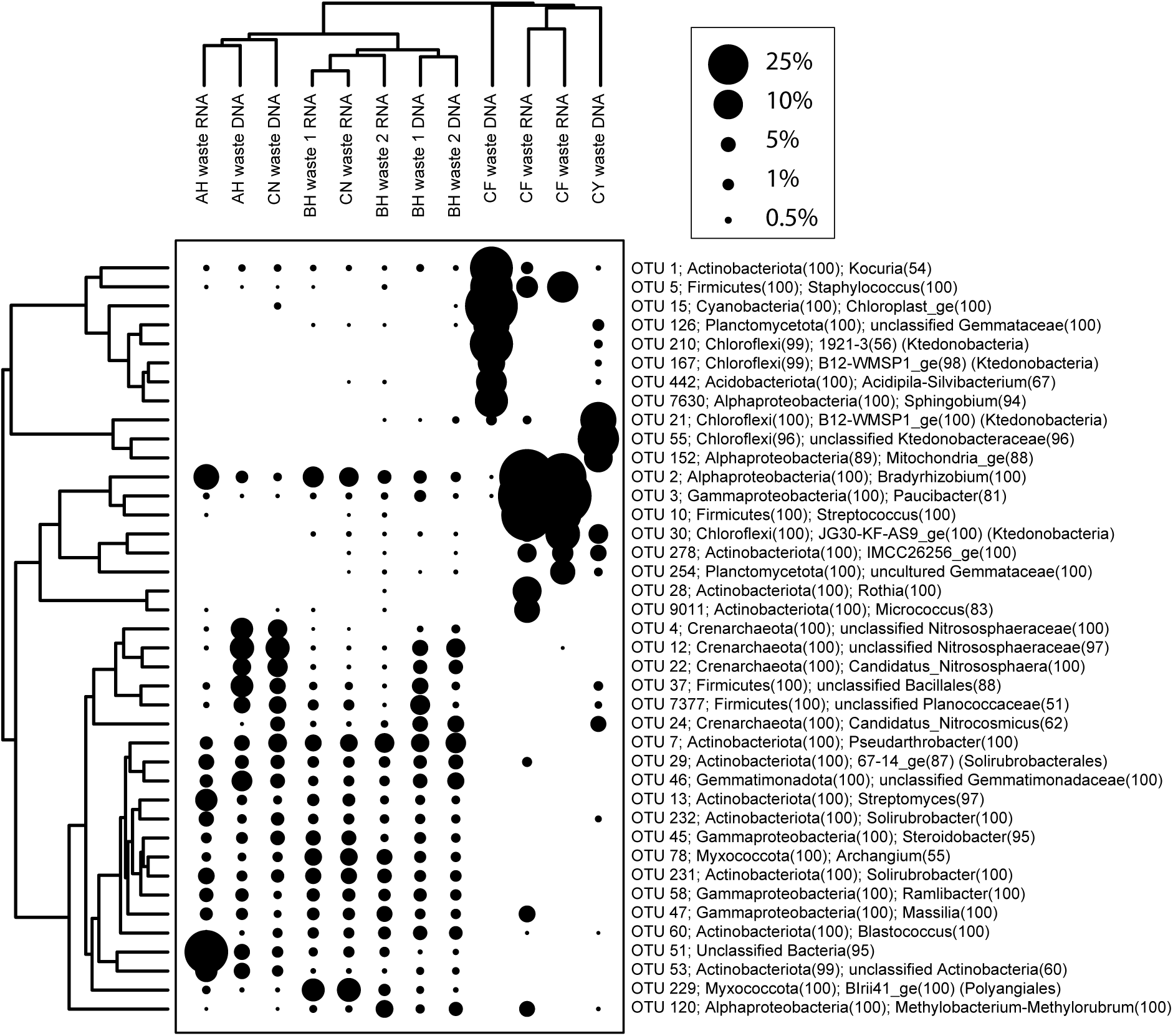
A two-way cluster analysis comparing 16S rRNA gene and transcript libraries from bulk waste rock. The points are scaled according to relative abundance (%). The taxonomic affiliation shows phylum and genus for each OTU, except for those that unclassified at the genus level, for which the highest available taxonomic classification is provided. BH waste 1 and 2 refer to samples from the upper and lower bench, respectively, and the two CF waste RNA libraries are from separate composite samples from that site (Fig 6; Table S1).

### Cell counts

We performed cell counts on composite waste rock samples from sites AH, BH, CN, and CY, and for site CF, we counted four individual samples. Bulk waste rock had between 6.1×10^7^ and 1.1×10^10^ cells g^-1^, with the lowest cell abundance at site CF and the highest at site CN (Table 2). Overall, tailings had lower cell abundance, between 2.0×10^6^ and 6.9×10^8^ cells g^-1^ (Table 2). At sites CF and CN, cell counts were lowest in tailings sampled below the visible oxidation fronts, but were at a similar order of magnitude to samples from at and above the oxidation front (Table 2).

**TABLE 2.** Cell counts from select waste rock and tailings.

| **Sample** | **Location** | **Sample type** | **Cells/g wet sediment** | **Sample notes** |
| --- | --- | --- | --- | --- |
| **AH24-13** | Alhambra | bulk waste | $9.9 \times 10^8$ | soil cover, waste composite |
| **BH24-13** | Black Hawk | bulk waste | $1.6 \times 10^9$ | soil cover, waste composite |
| **BH24-23** | Black Hawk | bulk waste | $1.5 \times 10^9$ | soil cover, waste composite |
| **CN23-13** | Center | bulk waste | $1.1 \times 10^{10}$ | soil cover, waste composite |
| **CY23-13** | Carlisle | bulk waste | $3.9 \times 10^9$ | waste composite |
| **CF22-601\*** | Copper Flat | bulk waste | $2.4 \times 10^8$ | |
| **CF22-603\*** | Copper Flat | bulk waste | $1.2 \times 10^8$ | |
| **CF22-604\*** | Copper Flat | bulk waste | $6.1 \times 10^7$ | |
| **CF22-606\*** | Copper Flat | bulk waste | $4.7 \times 10^8$ | |
| **CF23-621** | Copper Flat | tailings | $1.0 \times 10^8$ | above oxidation front |
| **CF23-622** | Copper Flat | tailings | $9.0 \times 10^7$ | at oxidation front |
| **CF23-623** | Copper Flat | tailings | $2.0 \times 10^7$ | below oxidation front |
| **CF23-631** | Copper Flat | tailings | $4.8 \times 10^7$ | above oxidation front, tailings composite |
| **CY23-20** | Carlisle | tailings | $9.7 \times 10^7$ | above oxidation front |
| **CY23-21** | Carlisle | tailings | $6.9 \times 10^8$ | at oxidation front |
| **CY23-22** | Carlisle | tailings | $1.6 \times 10^8$ | below oxidation front |

\*Individual (composite sample was not available for cell counting from the CF composite)

## DISCUSSION

### Different waste types have distinct microbial communities

Overall, microbial communities varied strongly among the different waste types (seeps and adits, bulk waste rock, and tailings). The acidic seeps and adits had the least diverse communities, with abundant taxa of known acidophilic iron- and sulfur-oxidizing microorganisms that are common in acid rock drainage, such as *Leptospirillum* spp., *Sulfuritalea* spp., and *Acidiphilium* spp., *Cuniculiplasma* spp., *Acidibacter* spp., and *Ferrithrix* spp. (24, 26, 27, 31, 100). There are also photosynthetic primary producers present, based on chloroplasts from the unicellular algal taxa *Chlamydomonas* and *Euglena*. Overall, the most abundant organisms in the acidic seeps and adits only occur at low levels in the waste rock and tailings (Fig. S2).

In contrast, microbial communities in the bulk waste rock and tailings were more diverse, and were dominated by different microorganisms, including abundant taxa from genera and families without cultured representatives (Fig. 3, S3). Many of the OTUs identified in the bulk waste and tailings that were classified to the genus level are more commonly associated with soil or other non-mining environments. For example, based on rRNA gene and transcript libraries, *Solirubrobacter* spp. and *Kocuria* spp. are abundant and active in the bulk waste piles sampled here, and are commonly found in desert soils that have been found to be highly UV-C resistant and dessication resistant (101–103). Bulk waste also contains unclassified members of the *Ktedonobacteraceae* (Fig. S3), which has been found to have a degree of heavy metal resistance and are often associated with metal-impacted soils (104, 105). Waste rock communities also include members of the Crenarchaeota and Thaumarchaeota such as *Candidatus* Nitrososphaera, *Candidatus* Nitrocosmicus, and *Candidatus* Nitrosotalea in rRNA gene and transcript libraries. These are three groups of ammonia-oxidizing archaea (AOA) that have been found in a variety of environments including ARD (106), wastewater treatment plants (107), and, more commonly, soils (108–113). There are other genera of common soil nitrogen cyclers present in our bulk waste sample microbial communities including *Noviherbaspirillum* (114, 115), *Bradyrhizobium* (116, 117), and *Nitrospira* spp. (111, 118). Some of the taxa that are abundant in tailings, but not seeps or the bulk waste, also include organisms common to soils. For example, *Ramlibacter* spp. have been observed in soils and associated with plant rhizomes (119–122). *Sphingomonas* spp. have been found in a wide variety of environments including freshwater (123) and soils (124, 125), including reclaimed amended tailings piles (87), and *Enhydrobacter* sp. has been identified in metal-impacted soils specifically soils containing high concentrations of cadmium (Cd) (126, 127).

The waste rock communities at sites AH, BH, and CN had both the highest diversity and species richness compared to CF and CY (Fig. 2). This is likely because the waste rock at AH, BH, and CN had 10-15 cm of soil cover over the top of the waste rock piles, while waste rock and tailings piles at CF and CY had no such cover. A cover provides additional opportunities for microbial colonization from the overlying soil, and increases nutrient availability and substrate attachment opportunities. There has been limited work assessing the microbial communities of bulk waste rock, both with and without a soil cover. Our results are consistent with work by (128) which found that bulk waste rock that was covered with a soil overburden had significant microbial contributions from the soil cover, and that samples close to the soil-waste interface were more diverse than samples from deeper in the waste rock pile. In our samples, the soil cover appears to have created a more homogenous microbial environment, as libraries from individual CF and CY samples are much more variable compared to individual samples from AH, BH, and CN (Fig. 2, 4).

In this study, we focused on the shallow surface of the waste rock and tailings. Chen et al. (83) found that *Proteobacteria* were primarily dominant in the upper layers of the tailings pile, and that the community shifted at depth to contain more *Firmicutes* and *Desulfobacterota*. This community change was attributed to geochemical variables such as pore-water pH increases with depth and dissolved metal ion concentrations, most notably iron. Other studies have suggested that community changes with depth also hinge on the disappearance of oxides and oxygen, leading to an increase in sulfate- and iron-reducing microorganisms (53, 88, 129). The only areas where we sampled with depth were in tailings piles at sites CF and CN, where we were able to sample above, below, and at a visible redox interface (Fig. 1a, S1). *Acidibacter* spp., which is a genus of iron-reducing bacteria (130), are present at and below the redox interface at CY, but otherwise we don’t see any clear trends in the specific phyla identified in (83) or any abundant known sulfate or metal reducers (Fig. S8). However, many libraries contain OTUs from genera, families, and higher taxonomic groupings, so it is possible that these samples contain unrecognized sulfate or iron reducers.

Overall, ordination analyses showed that differences among microbial communities correspond to elemental content and other geochemical variables among the sites (Fig. 5). In addition, some OTUs are statistically significantly correlated with geochemical variables such as percent sulfur, iron, and other metals that could indicate a response to rock-associated chemical energy resources or toxic compounds (Fig. S9). For example, OTUs corresponding to “*Candidatus* Nitrososphaera” and the group KD4-96 in the *Chloroflexi*ota are abundant in samples with high iron and low sulfur concentrations that have low copper and lead (Fig. S9), and uncultured members of order Vicinamibacterales (phylum *Acidobacteriota*) and *Gaiellales* (*Actinobacteriota*) are similarly negatively correlated with sulfur, copper, and lead content (Fig. S9), seemingly independent of site. Perhaps these taxa are sensitive to metals. In contrast, there are not any abundant OTUs that show clear correlations with increasing sulfur content. Some taxa from samples with the highest sulfur include *Dechloromonas*, *Rothia*, *Corynebacterium*, *Enhydrobacter*, *Sphingomonas*, which are not genera typically associated with inorganic sulfur compound oxidation, but these taxa are only abundant in a few high sulfur samples. There are some OTUs that are positively correlated with rare earth element content, such as OTUs classified Archangium (*Myxococcota*) and TRA3−20 in the *Gammaproteobacteria* (Fig. S9), although these taxa could instead be responding to other factors, such as high concentrations of other elements like As or Co that are also abundant in these samples (Table S1). OTUs classified as IMCC26256 (*Actinobacteriota*) and B12-WMSP1 (*Ktedonobacteria*, *Chloroflexi*ota) are abundant in rRNA gene and transcript libraries from samples with high Cu and low pH from uncovered waste rock at CY and CF (Fig. S9), which could indicate an association with copper sulfide minerals at these sites. However, future meta-omic or culture-based analyses will be necessary to evaluate the genomic and metal-cycling capabilities of these and other uncultured and unclassified organisms associated with these historic wastes.

### Implications for microbiological characterization of historic waste rock and tailings

Waste rock and tailings can be challenging environments for microbiological analysis due to their heterogeneity, metal content, and significant spatial extent on the order of 10s of meters up to kilometers. To attempt to capture this heterogeneity, we compared individual versus composite samples, and showed that the composite samples were effective at representing the heterogeneity of the waste pile microbial communities for the waste piles sampled here (Fig. 6). On one hand, collecting and analyzing multiple samples from each waste rock pile provided an opportunity to draw more careful correlations between geochemical variables and help parse out site specific effects (Fig. 5, S9). However, on the other hand, a large number of samples for microbiological analysis can be cost prohibitive, so collecting a composite sample is a reasonable approach for characterizing the waste-associated microbial communities from large waste rock and tailings piles, at least for piles that are on the order of several 10s of meters like those sampled here.

### Implications for mine waste biogeochemistry and metal mobilization

Microbial communities associated with acid mine drainage and acidic, sulfide mineral-rich ores often have abundant iron- and sulfur-oxidizing populations. These organisms are important for metal release in biomining operations. Iron-oxidizers serve as “oxidant generators” that produce Fe(III), which is the primary oxidant for metal sulfide minerals; sulfur oxidizers are “acid generators” that produce sulfuric acid and contribute to the breakdown of some sulfide minerals; and heterotrophs act as “janitors” that remove low molecular weight organic carbon compounds that are often toxic to the autotrophic Fe and S oxidizing populations (131, 132). While we found this type of community in the acidic seeps and adits, the waste rock and tailings had a very different structure that was dominated by taxa more common to soil and that are associated with the oxidation of organic carbon and inorganic nitrogen compounds. Some taxa in the waste rock and tailings are known to oxidize or reduce iron and sulfur compounds, such as *Sulfurifustis* (133) and *Acidibacter* (130) (Fig. 3), but these were not very abundant. This differs from previous work on circumneutral mine waste, especially work utilizing culture-dependent methods, where a higher number of acidophilic and neutrophilic iron- and sulfur-oxidizing microorganisms have been identified (73, 78, 80, 85, 86, 88, 89, 134). It is likely that the communities within the waste rock piles sample here are not mobilizing metals to the same extent we would see in ARD and bioleaching heaps.

The presence of a soil-like microbial community may indicate successful reclamation. The revegetation of tailings piles has been shown to change microbial community assemblage in favor of nitrogen-cycling and rhizome-associated microorganisms (81), thereby halting metal mobilization and preventing the formation of ARD (41). For example, prior studies have found that reclaimed vegetated tailings piles contain high abundances of *Nitrospira* spp. and *Firmicutes*. (79, 81, 84). In semi-arid areas in particular, microorganisms common to soil microbiomes were found in high abundance on abandoned tailings piles, and 47% of the taxa present in these communities were associated with drought and UV stress conditions (87), like the *Solirubrobacter* spp. and *Kocuria* spp. observed here.

With critical minerals in high demand and the push to identify and quantify non-traditional metal resources, waste rock and tailings represent a substantial resource that could theoretically be accessed using in situ leaching or other biomining techniques. However, we don’t know if the more abundant sulfur- and metal-cycling organisms such as *Sulfurifustis* and *Acidibacter*, or some of the rare populations like *Acidiphilium*, could be stimulated to mobilize metals and critical minerals in these deposits. Changing geochemical conditions to favor these organisms, such as by increasing the temperature or moisture conditions or adding elemental sulfur or ferrous iron to favor chemolithoautotrophic microbial growth, could stimulate leaching (55, 65, 134–136). In addition, like other historic mine waste resources (83), these deposits have many unknown populations that may be mobilizing or immobilizing metals via metabolic pathways that are not yet well-understood or recognized. A more complete understanding of the microbial assemblages that develop *in situ* may provide new insight into the most appropriate organisms and processes for metal recovery and recycling from these and other historic mine waste deposits.

## MATERIALS AND METHODS

### Field sites and sampling procedures

Samples were collected from five inactive mine sites: Copper Flat mine (Sierra County), Carlisle mine, Center mine, Alhambra mine, and Black Hawk mine (Grant County), abbreviated CF, CY, CN, AH, and BH, respectively. Samples of all available waste types were collected at each site, and included acidic seeps, flooded adits, bulk waste rock, and tailings (Table 1).

For waste rock sampling, material for geochemical analyses was sampled from the top 15 cm as described in (90), (91), and (9). Samples for microbiological analysis were collected by using sterile implements to expose a fresh surface and scrape approximately 5 mL of material from these same holes into a sterile 15 or 50 mL tube. Samples were mixed by shaking, and approximately 500µL was removed into a 2 mL microcentrifuge tube for cell counting. The larger sample was then immediately frozen on dry ice for nucleic acid extraction and stored at -80°C upon return to the NMT Geobiology Lab. The smaller subset was fixed with paraformaldehyde (PFA) as in (137) and stored at 4°C. For large waste rock piles, samples were collected from six surface sites selected randomly around the perimeter in an attempt to evaluate and capture the heterogeneity of the large pile. Samples were preserved individually, along with a composite sample that was prepared by combining approximately 5 mL from each of the six locations and preserving it as for the individual samples. Four sampling locations had a soil cover: the tailings at CF, and bulk waste rock at CN, BH, and AH. Where a soil cover was present, the hole was dug so that waste rock material was sampled and not the overlying soil cover.

Because of the larger size of the tailings pile at CF, individual and composite samples were collected as for bulk rock above. At sites CF and CN, tailings displayed oxidation fronts that were clearly visible by color changes, and were sampled above, at, and below these fronts. Sediments of seep and adit sediments were collected from the top 0.5 cm and preserved as described above.

### Nucleic acid extraction and rRNA gene and transcript library preparation

DNA was extracted using a DNeasy PowerSoil Pro DNA isolation kit (Qiagen, Hilden, Germany) following the standard kit protocol with one modification: samples were subject to bead beating for 20, 40, and 60 seconds at 2500 RPM using a Mini-Beadbeater (Biospec Products, Bartlesville, OK, USA), and a 250 µL aliquot was removed at each time point as in (99). To avoid the “nugget effect”, each sample was extracted at least twice. The V4 region of the 16S rRNA gene (DNA) was amplified using 10mM V4 Nextera primers (Integrated DNA Technologies, Coralville, IA, USA; forward primer tail TCGTCGGCAGCGTCAGATGTGTATAAGAGACAG; reverse primer tail GTCTCGTGGGCTCGGAGATGTGTATAAGAGACAG). PCR was performed with the AllTaq PCR Core Kit (Qiagen) with initial denaturation for 5 minutes at 94°C followed by 30 or 35 cycles of 94°C for 30s, 52°C for 45s, and 72°C for 30s, with a final elongation at 72°C for 10 minutes. Samples were then held within the thermocycler at 4°C until they were moved to a -20°C freezer. Samples were run for 30 cycles initially, and if no amplification was observed, the sample was re-amplified for 35 cycles. A DNA extraction blank was also subjected to PCR to check for contamination during the DNA extraction process, and a PCR reagent blank was amplified to check for PCR reagent contamination. Extractions with visible PCR produce in blanks were not processed further. We also tested DNeasy PowerBiofilm Kit (Qiagen) and DNeasy PowerSoil Kit (Qiagen), but had overall better performance with the DNeasy PowerSoil Pro kit (Qiagen).

RNA extraction was performed using the RNeasy PowerBiofilm RNA isolation kit (Qiagen), using the same bead-beating protocol as for DNA extractions, above, with a 150 µL aliquot removed after 20, 40, and 60 s of bead beating. DNA was removed using two DNase treatments, an on-column DNase treatment using DNase I (Qiagen) that was provided in the RNeasy PowerBiofilm RNA isolation kit, and any remaining DNA in the sample was digested using the Invitrogen TURBO DNA-free kit kit (Thermo Fisher Scientific, Waltham, MA, USA) and subsequently cleaned with the RNA Clean and Concentrator-5 kit (Zymo Research, Irvine, CA, USA). Reverse-transcription PCR was conducted using a OneStep *Ahead* RT-PCR Kit (Qiagen) with the same V4 primers as described above under the following program: 50°C for 10 minutes, followed by an initial denaturation for 5 minutes at 94°C, and then 30 or 35 cycles of 94°C for 10s, 50°C for 10 s, and 72°C for 15 s before finishing with 5 minutes at 72°C and holding at 12°C until the PCR products were moved to the -20°C freezer for storage. Samples were run for 30 PCR cycles initially, and were re-run at 35 cycles if no RNA amplification was observed. To ensure that samples did not contain residual DNA, we amplified the RNA extract according to the DNA PCR specifications described above. If DNA amplification was observed after 35 cycles, the Invitrogen TURBO DNA-free (Qiagen) treatment and RNA Clean and Concentrator (Zymo) process was repeated until no amplification was observed (usually no more than one additional treatment was necessary). A PCR reagent blank was also included in each PCR run to check for PCR reagent contamination.

PCR products were barcoded and sequenced at the University of Minnesota Core Genomics Center (UMGC) on an Illumina MiSeq with version 3 chemistry and 300 paired end cycles. DNA extraction blanks and PCR reagent blanks were sequenced alongside samples, even though no PCR products were visible.

### Cell counting

In order to detach cells from mineral and soil particle surfaces, a small amount of the PFA-fixed sample and sonicated using a FisherBrand Model CL-18 dismembrator (Fisher Scientific, Hampton, NH, USA) for 45 seconds on 100% amplitude. Cells were then stained with DAPI (4′,6-diamidino-2-phenylindole) (Biotium, Fremont, CA, USA) and filtered onto a 0.2 µM membrane filter (Cytiva Whatman, Marlborough, MA, USA). The filter was mounted on a glass slide, covered using Vectashield antifade mounting medium (Vector Laboratories, Newark, CA, USA), and observed under an Olympus BX63F compound epifluorescence microscope (Olympus Life Science, Waltham, MA, USA). Cells were counted from 10 randomly-selected grid sections on the slide, averaged, and converted to cells g^-1^ sediment based on the mass of sediment sonicated and accounting for the dilution at each step.

### Bioinformatic and statistical analyses

Quality filtering, trimming, and OTU calling followed the same procedures as in (137) and described below. Raw sequences were trimmed and filtered using Sickle (https://github.com/najoshi/sickle) to an average quality score of >28 (5’ trimming only) and a minimum length of 100 bp. Cutadapt v. 4.2 (138) was used to trim residual adaptors, and PEAR (138) was used to assemble the forward (R1) and reverse (R2) reads. Primer sequences were removed with prinseq v.0.20.4 (140). OTU calling (97% similarity) and chimera removal also followed procedures in (137) using USEARCH v. 11 (141) and VSEARCH v. 2.21 (142). Representative OTUs were determined with mothur v. 1.47 (143) and SILVA v. 138.1 (144) with a confidence cut-off score of 50. Following quality filtering and trimming, libraries with <10,000 reads were excluded from subsequent community analyses.

R Statistical Software RStudio v. 4.2.1 (145) was used to conduct statistical analyses. Additional R packages used included vegan (v. 2.6-4) (145), cluster (2.1.4) (147), ggplot2 (v. 3.5.1) (148). Rare OTUs (<0.01%) were excluded, and the remaining data was normalized using the angular transformation to deemphasize abundant OTUs. Non-metric multidimensional scaling (NMDS) ordinations were calculated using the *metaMDS()* function with four dimensions (k=4). ANOSIM and PERMANOVA analyses were calculated with the vegan package using default parameters (Bray-Curtis distance, 999 permutations, with *anosim()* and *adonis2()* functions). The statistical significance of comparisons between 3 or more groups was assessed with a pairwise analysis using the *pairwise.adonis()* function (149). Diversity was calculated using richness and the Shannon-Wiener Index, which takes into account the number of OTUs and their evenness, using functions *rarefy()* and *diversity()* after subsampling the OTU matrix to 10,000 reads. Stacked bar charts of OTU relative abundance were created using code from https://jkzorz.github.io/2019/06/05/stacked-bar-plots.html and color palettes for the stacked bar charts from Paletteer (https://github.com/EmilHvitfeldt/paletteer).

## Data availability

Raw sequences are available on the NCBI SRA under BioProject PRJNA1226971.

## ACKNOWLEDGMENTS

We would like to thank researchers and students from the economic geology research group at the New Mexico Bureau of Geology and Mineral Resources for their help with field work and site access including Bonnie Frey, Evan Owen, Mark Leo-Russell, Stellah Cherotich, Abena Acheampong-Mensah, Kyle Stafford, and Bob and Jakob Newcomer. We also thank Robert Seal and Kate Campbell from the USGS for assistance with field work and insightful discussion. We thank Jimmy Swift for his help with cell counts and microscopy. Site access was provided by Themac Resources and Santa Fe Gold Corp.

This work was supported by a USGS Earth MRI grant award #G22AC00510, and a Grant-in-Aid from the New Mexico Geological Society.

