## Supplemental Figures for "Historic mine waste contains diverse microbial communities that reflect waste type and geochemistry"

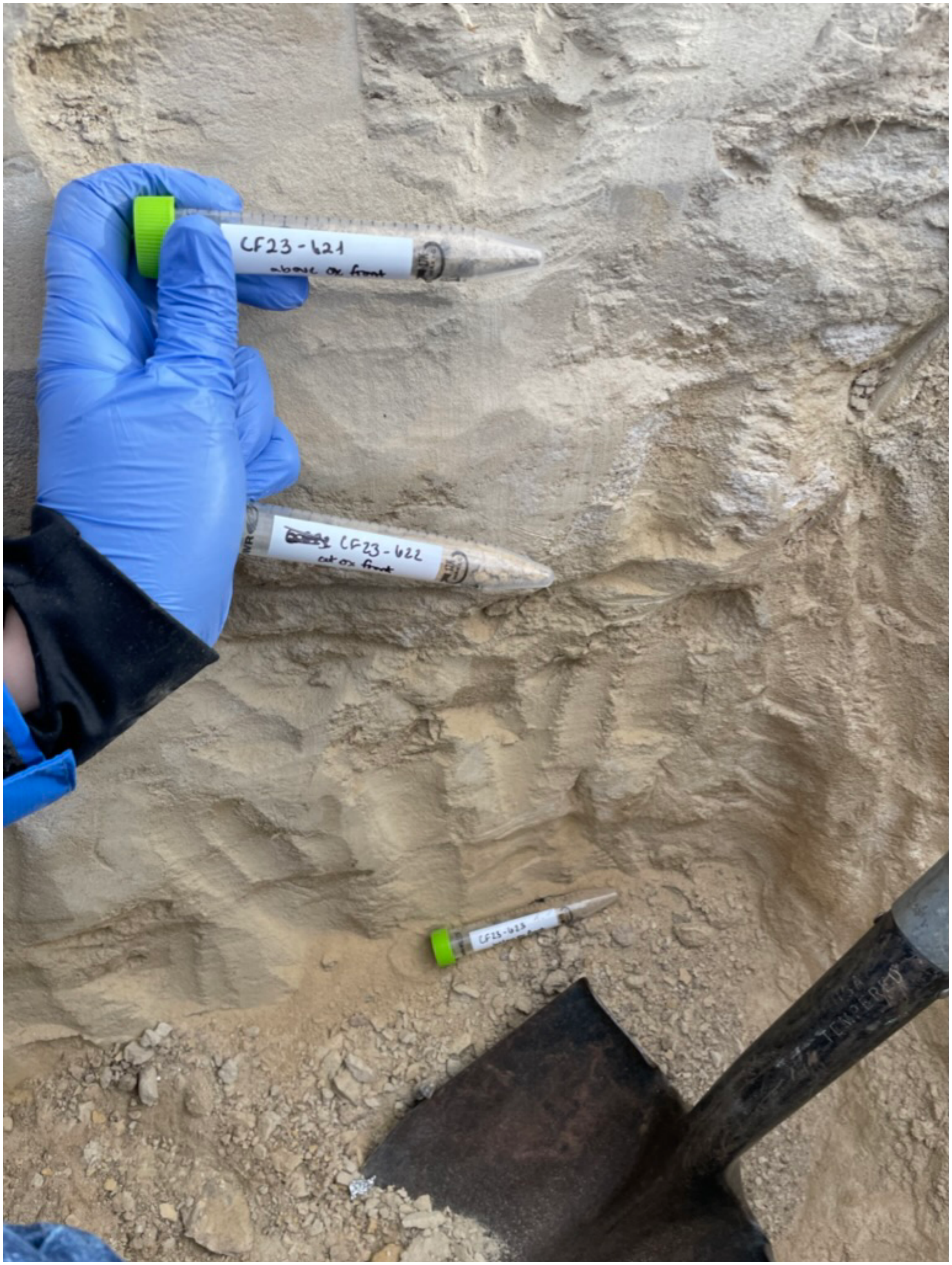


**SUPPLEMENTARY FIG S1**. Tailings samples collected from a 1 m pit at Copper Flat Mine. Samples were collected from above, at, and below the oxidation front, which is visible due to color changes from white, to beige, to yellow.

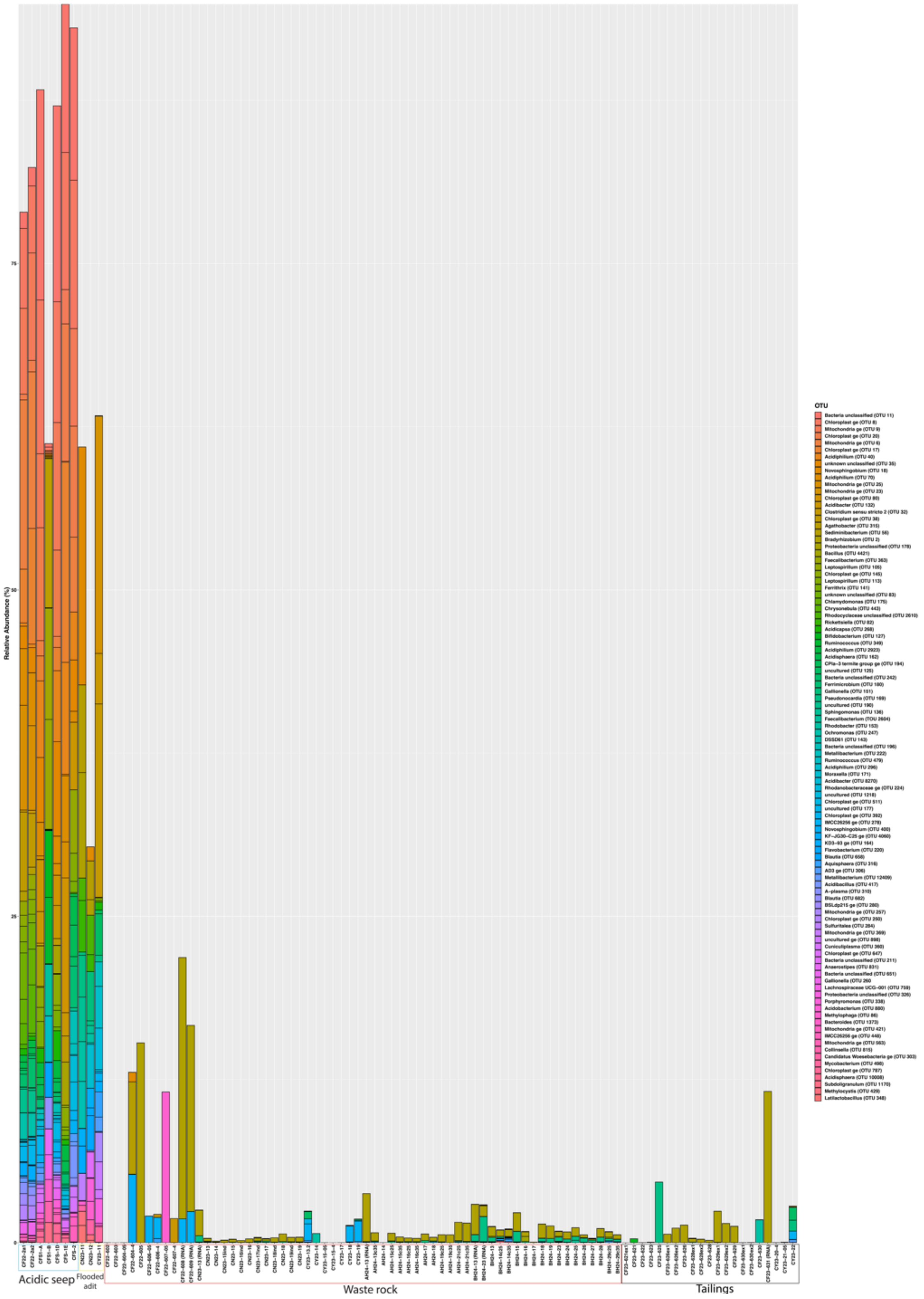


**SUPPLEMENTARY FIG S2.** Genus-level microbial community composition of the 100 most abundant OTUs in the acidic seep and adit samples. Bars representing samples are grouped by sample type (L to R): seep, adit, bulk waste rock, and tailings.

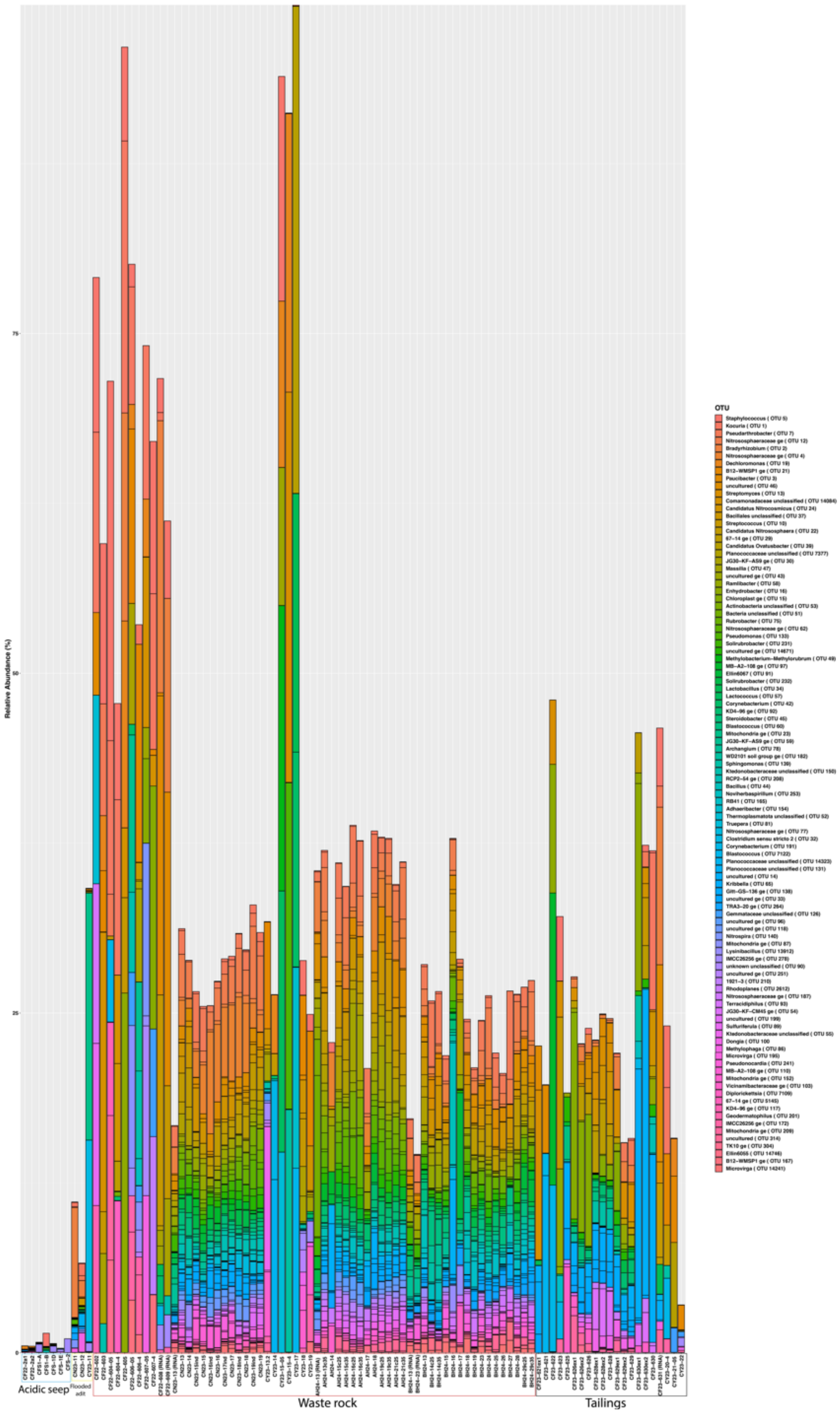


**SUPPLEMENTARY FIG S3.** Genus-level microbial community composition of the 100 most abundant OTUs in the bulk waste rock samples. Bars representing samples are grouped by sample type (L to R): seep, adit, bulk waste rock, and tailings.

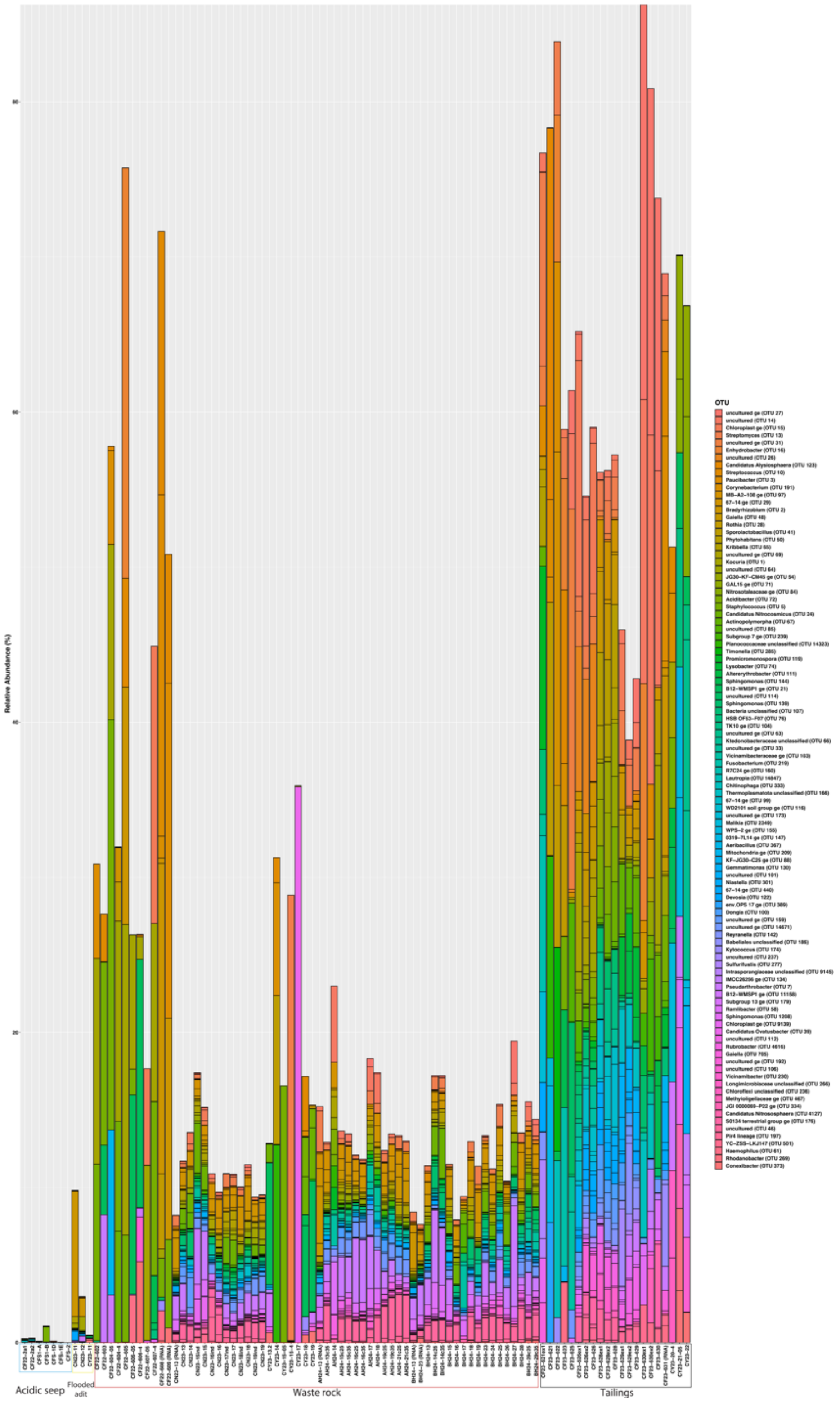


**SUPPLEMENTARY FIG S4.** Genus-level microbial community composition of the 100 most abundant OTUs in the tailings samples. Bars representing samples are grouped by sample type (L to R): seep, adit, bulk waste rock, and tailings.

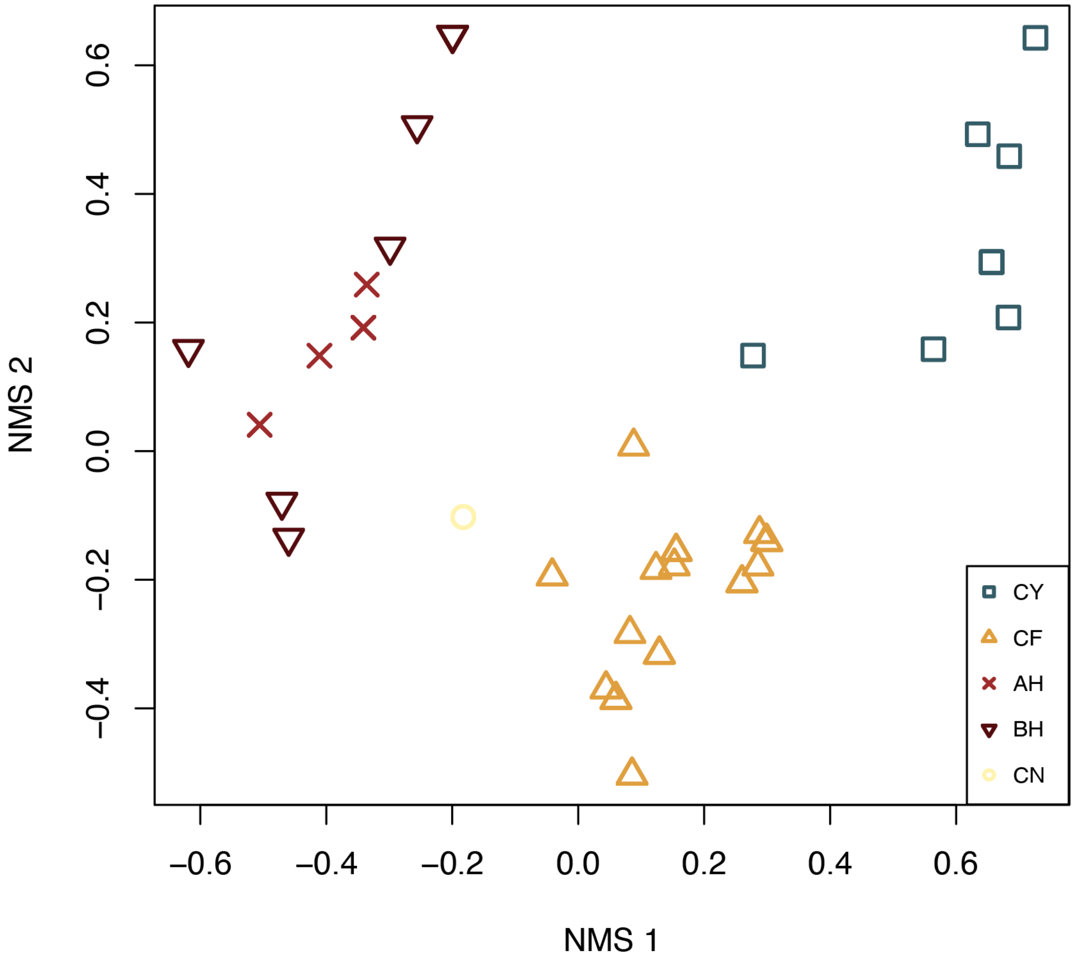


**SUPPLEMENTARY FIG S5**. Non-metric multidimensional scaling (NMDS) ordination of individual mine sites based on geochemical parameters alone. Parameters used to ordinate samples are carbon (%), sulfur (%), gold, silver, arsenic, cobalt, copper, iron (%), lead, total rare earth elements (TREE), uranium, and silica (%) (Supplementary Table S1).

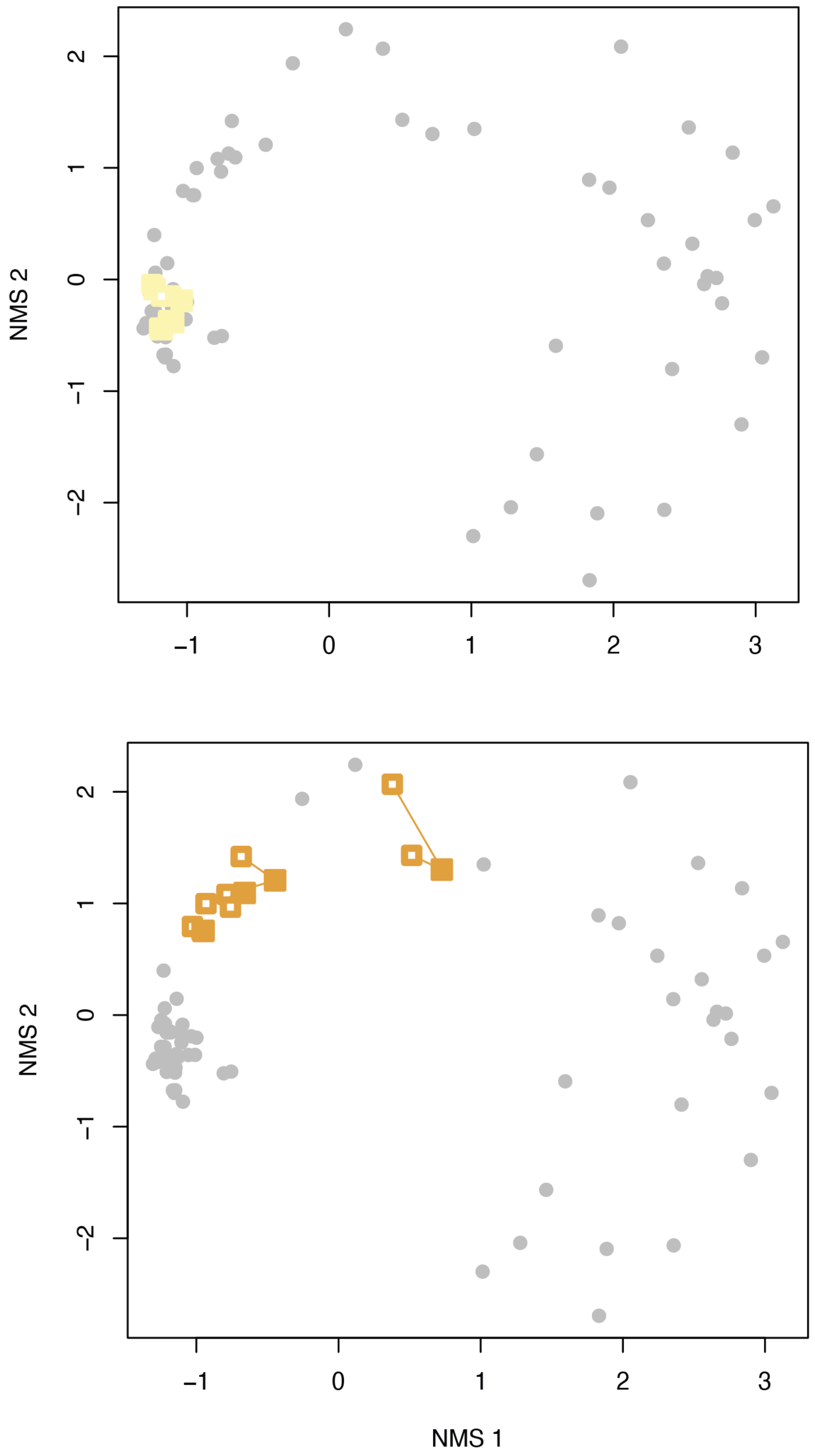


**SUPPLEMENTARY FIG S6**. A non-metric multidimensional scale (NMDS) ordination of samples from Center Mine (top, yellow) and Copper Flat Mine (bottom, orange) showing the similarity between individual DNA extracts (open squares) and the combined DNA extract (closed squares).

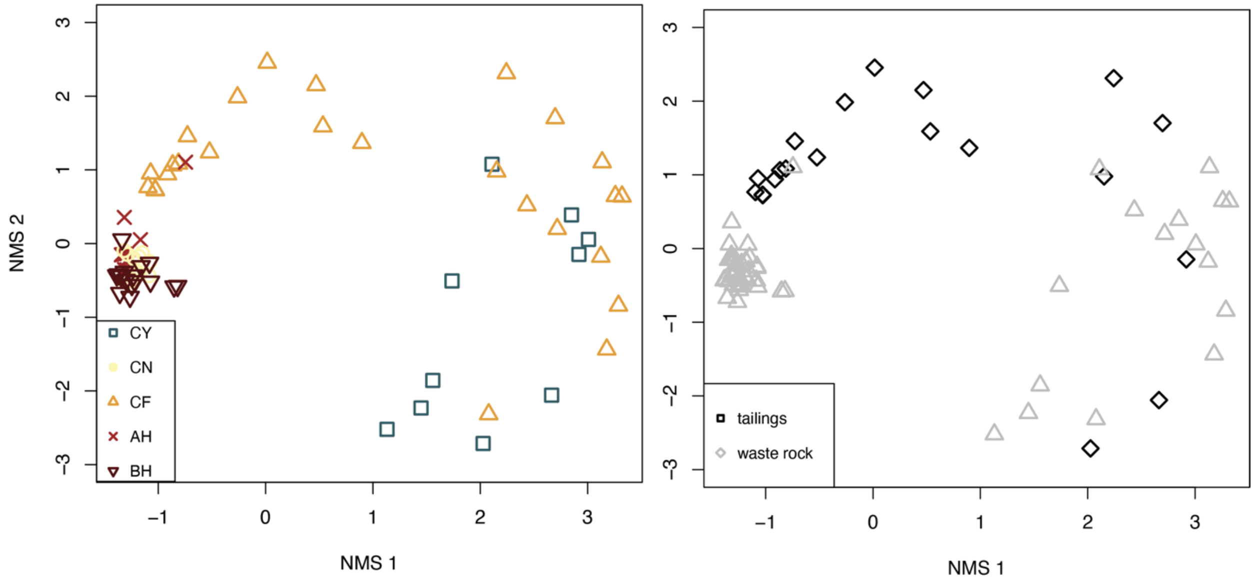


**SUPPLEMENTARY FIG S7**. Non-metric multidimensional scaling (NMDS) ordination of rRNA gene libraries only (no rRNA transcript libraries like in Figure 4 in the main text). Left: Samples are coded by location with samples from Center Mine as open yellow circles, Copper Flat Mine samples represented by open orange triangles, Alhambra Mine samples as red x's, Black Hawk Mine as dark red open upside-down triangles, and Carlisle Mine as open blue squares. Right: Samples are coded by sample type where tailings samples are represented by open black squares and waste rock samples are shown with open grey triangles.

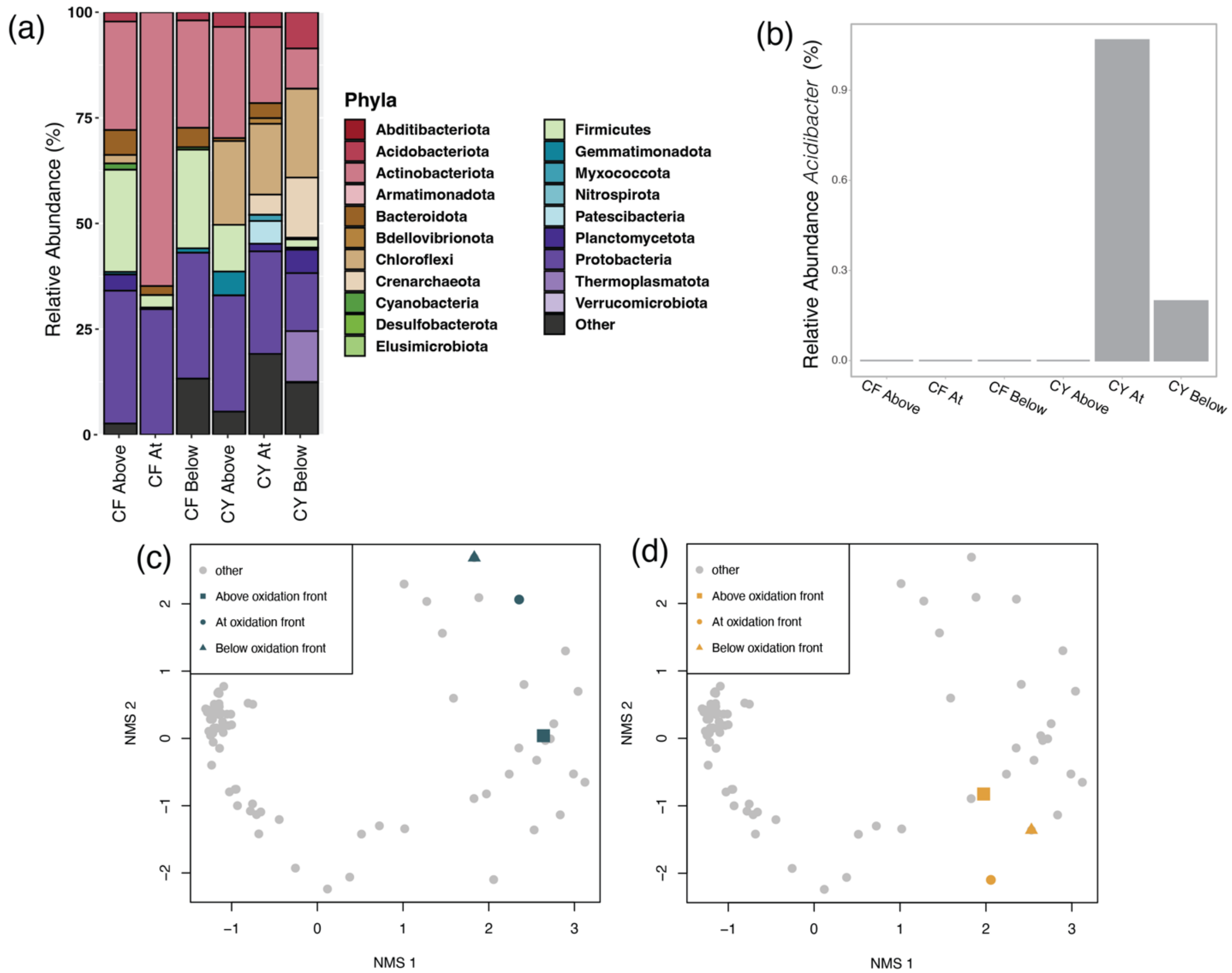


**SUPPLEMENTARY FIG S8**. (a) Phyla-level community composition of samples collected from Copper Flat (CF) and Carlisle Mines (CY) above, at, and below the visible redox interface (Fig. S1, S2). (b) The relative abundance of *Acidibacter* spp. across the redox interface at CF and CY. (c) NMDS ordination comparing microbial communities across the redox interface at CY. (d) NMDS ordination comparing microbial community composition across the redox interface at CF.

**SUPPLEMENTARY FIG S9 (next two page)**. Correlations between select OTUs and geochemical parameters. The text at the top of the figure indicates Pearson's correlation coefficient and statistical significance (R and p-value) for the relationship shown. The taxonomic affiliation for the OTUs includes phylum- and genus-level classifications, or the highest available taxonomic classification for uncultured taxa. Confidence scores >50 are provided in parentheses.

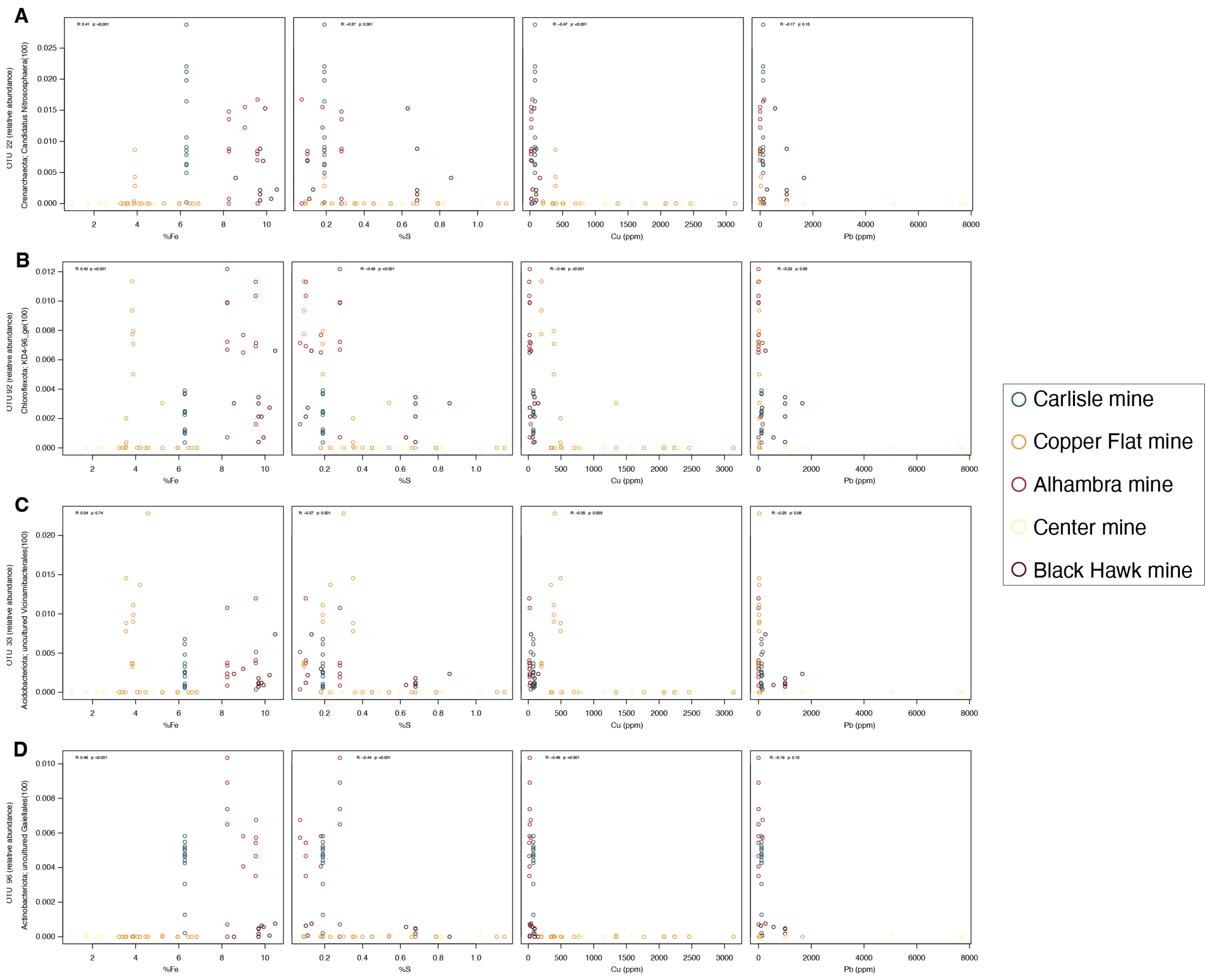
A
B
C
D
OTU 22 (relative abundance)
Crenarchaeota; Candidatus Nitrososphaera(100)
OTU 92 (relative abundance)
Chloroflexota; KD4-96_ge(100)
OTU 33 (relative abundance)
Acidobacteriota; uncultured Vicinamibacterales(100)
OTU 96 (relative abundance)
Actinobacteriota; uncultured Gaiellales(100)
%Fe
%S
Cu (ppm)
Pb (ppm)
Carlisle mine
Copper Flat mine
Alhambra mine
Center mine
Black Hawk mine

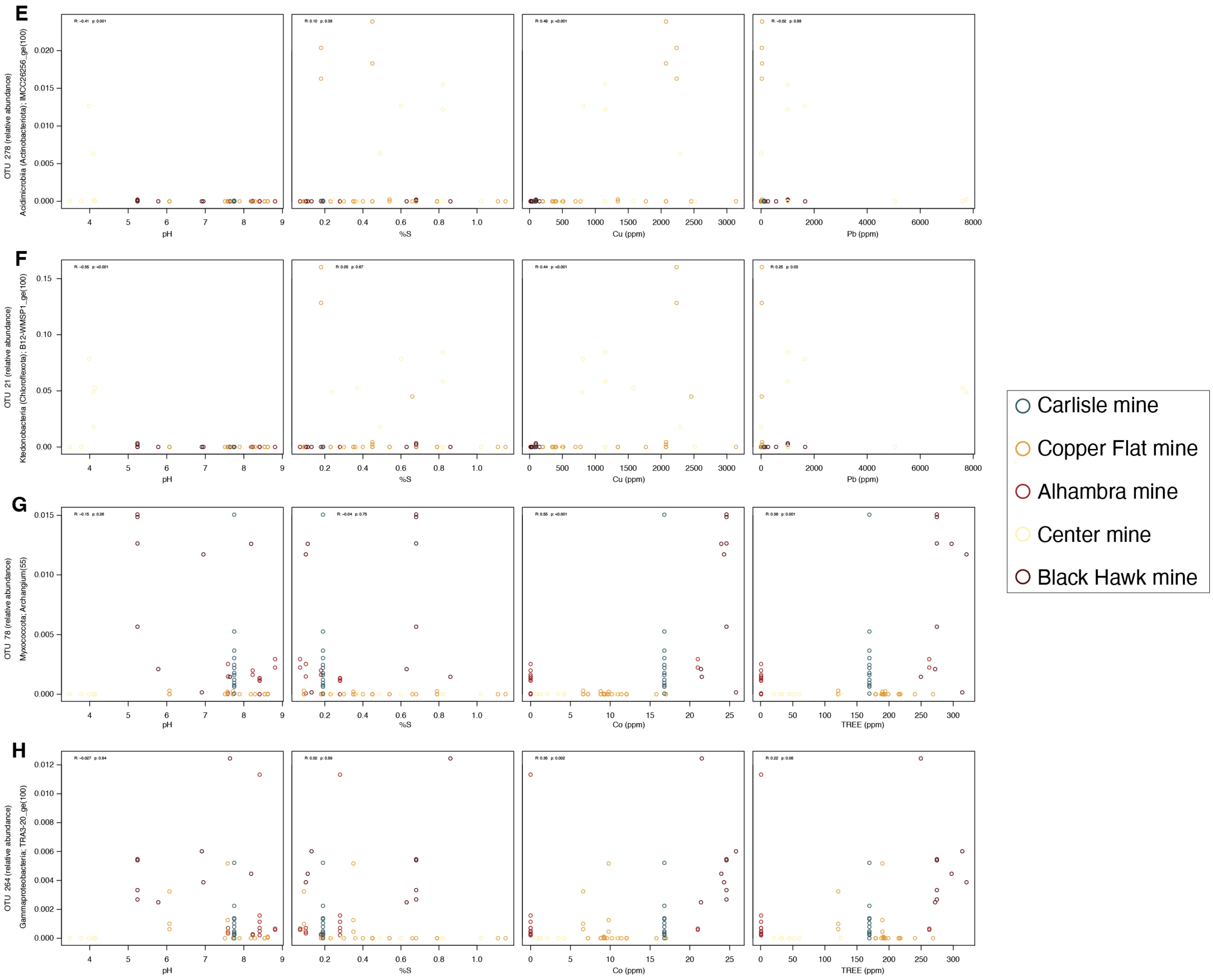
E
F
G
H
OTU 278 (relative abundance)
Acidimicrobiia (Actinobacteriota); IMCC26256_ge(100)
OTU 21 (relative abundance)
Ktedonobacteria (Chloroflexota); B12-WMSP1_ge(100)
OTU 78 (relative abundance)
Myxococcota; Archangium(55)
OTU 264 (relative abundance)
Gammaproteobacteria; TRA3-20_ge(100)
pH
%S
Cu (ppm)
Pb (ppm)
Co (ppm)
TREE (ppm)
Carlisle mine
Copper Flat mine
Alhambra mine
Center mine
Black Hawk mine
